# Control of epiblast cell fate by mechanical cues

**DOI:** 10.1101/2024.06.24.600402

**Authors:** Charlène Guillot, Yannis Djeffal, Mattia Serra, Olivier Pourquié

**Affiliations:** Institute of Genetics Reproduction and Development (iGReD) University of Clermont Auvergne, CNRS UMR6293, INSERM U1103, 63000 Clermont-Ferrand, France; Department of Genetics, Harvard Medical School, Boston, Massachusetts, 02115, USA; Department of Pathology, Brigham and Women’s hospital, Boston, Massachusetts, 02115, USA; Physics Department, UC San Diego, La Jolla, California, 92093, USA; Mondor Institute for Biomedical Research (IMRB) Paris Est University, INSERM U955-E10, 94010 Créteil, France

## Abstract

In amniotes, embryonic tissues originate from multipotent epiblast cells, arranged in a mosaic of presumptive territories. How these domains fated to specific lineages become segregated during body formation remains poorly understood. Using single cell RNA sequencing analysis and lineage tracing in the chicken embryo, we identify epiblast cells contributing descendants to the neural tube, somites and lateral plate after completion of gastrulation. We show that intercalation after cell division generates important movements of epiblast cells which lead to their relocation to different presumptive territories, explaining this broad spectrum of fates. This tissue remodeling phase is transient, being soon followed by the establishment of boundaries restricting cell movements therefore defining the presumptive territories of the epiblast. Finally, we find that the epiblast faces distinct mechanical constraints along the antero-posterior axis, leading to cell fate alterations when challenged. Together, we demonstrate the critical role of mechanical cues in epiblast fate determination.

## Introduction

In amniotes such as chicken or mouse, most embryonic tissues derive from a superficial epithelial layer of pluripotent cells called epiblast ^1,2^. Cells from this tissue either become internalized in the primitive streak (PS) during gastrulation to form the endoderm and mesoderm or they remain superficial and give rise to ectoderm. Epiblast ingression starts in the PS which forms in the posterior region of the embryo and progressively extends anteriorly as gastrulation progresses. Gastrulation is often considered to last until full PS extension, when endodermal as well as heart, head mesoderm and notochord territories of the epiblast have ingressed. After it reached its maximal size, the PS begins to regress, laying in its wake the mesodermal territories of the trunk of the embryo which continue to ingress from the epiblast adjacent to the PS. This later phase of building the posterior part of the body is often considered to represent the continuation of gastrulation ^3–5^.

The fate of epiblast cells has been identified with fate mapping experiments where cells were marked to track their descendants. This defined the contribution of epiblast regions to specific embryonic territories and their final fates ^6–21^. Fate mapping the epiblast adjacent to the regressing PS in chicken and mouse embryos established that the domain anterior to the PS gives rise to the neural plate which forms the future anterior nervous system. The node, which marks the anterior tip of the PS, yields the prechordal plate and notochord. The anterior epiblast adjacent to the node and the anterior PS contains the neuro-mesodermal precursors (NMPs) which contribute both to paraxial mesoderm (PM) and spinal cord ^15,22,23^. More posteriorly along the PS lie the trunk mesoderm presumptive territories. These are organized along the AP axis of the epiblast flanking the PS according to their future medio-lateral location in the embryo (Supplementary Figure 1) ^7,8,14,18,24^. Thus, the territories of paraxial, intermediate, lateral plate and extraembryonic mesoderm are progressively found more posteriorly. Except for the NMPs, most of these territories appear to be specified to fates restricted to one single germ layer ^22,23^. During PS regression, these epiblast territories become progressively exhausted in a posterior to anterior direction until only the most anterior territory containing the NMPs and the node descendants (ie the notochord precursors) remains, contributing most of the tail bud progenitors (Supplementary Figure 1) ^22,25^.

Epiblast fate maps identified regions of overlap between the different mesodermal fates ^7^. Moreover, heterotopic grafting experiments as well as manipulating cell signaling demonstrated some level of plasticity of the epiblast ^15,26–29^. Whereas fate maps usually characterize the fate of cell populations they provide limited information on the lineage of individual cells. Thus, whether these overlapping regions arise from a mixed population of fated progenitors or from cells with broader developmental potential that can give rise to multiple mesodermal lineages remains undefined.

Here we use single cell RNA sequencing (scRNAseq) data to demonstrate that except for NMPs, mesodermal progenitors of the different presumptive territories of the epiblast do not exhibit significant transcriptome differences and thus constitute a remarkably homogeneous population. A distinct transcriptomic identity of the different mesodermal territories is only detected after their ingression in the PSM. We show that NMPs are maintained during axis elongation while the other epiblast progenitors which contribute to LP and extraembryonic mesoderm (EE) disappear before completion of PS regression. We performed heteropic and heterochronic grafting experiments to show that these two types of epiblasts exhibit significant plasticity during PS regression. We develop novel bioinformatic tools that predict a broader spectrum of cell fates for epiblast cells than demonstrated by classical fate maps. We validate these predictions using single cell fate tracking experiments to demonstrate the existence of epiblast precursors able to give rise to neural and PM cells also contributing to LP derivatives. Our experiments further suggest that acquisition of these different identities is possible due to significant epiblast cell movements triggered by cell intercalation after cell division. This mechanism ensures the dispersion of the descendants of epiblast cells and their relocation to distantly located presumptive territories. We also show that the anterior and posterior epiblast are exposed to very different mechanical environments. Altering the mechanical constraints of the epiblast can result in important cell fate changes in epiblast arguing that mechanical cues play an important role in the specification of epiblast cell fates.

## Results

### Limited heterogeneity of the transcriptome of epiblast cells

We generated a single cell RNA sequencing (scRNAseq) dataset of the PS and adjacent epiblast region at stage 4HH, 5HH, 6 somites and 35 somites. Using the inDrops pipeline ^30^, we generated single-cell transcriptomes of 11,812 cells from these 4 different embryonic stages. To identify cellular states, we developed a new unbiased clustering method based on differentially expressed genes, that can be used alongside known clustering methods such as Leiden or Louvain. This method, called Opticlust, considers the significance of the genetic profile of each identified cell population in our scRNAseq dataset (Figure 1A, left). Specifically, upon Leiden clustering, the resolution parameter is automatically determined such as all the clusters have a significant p-value for their first ranked differentially expressed genes.

**Figure 1:**
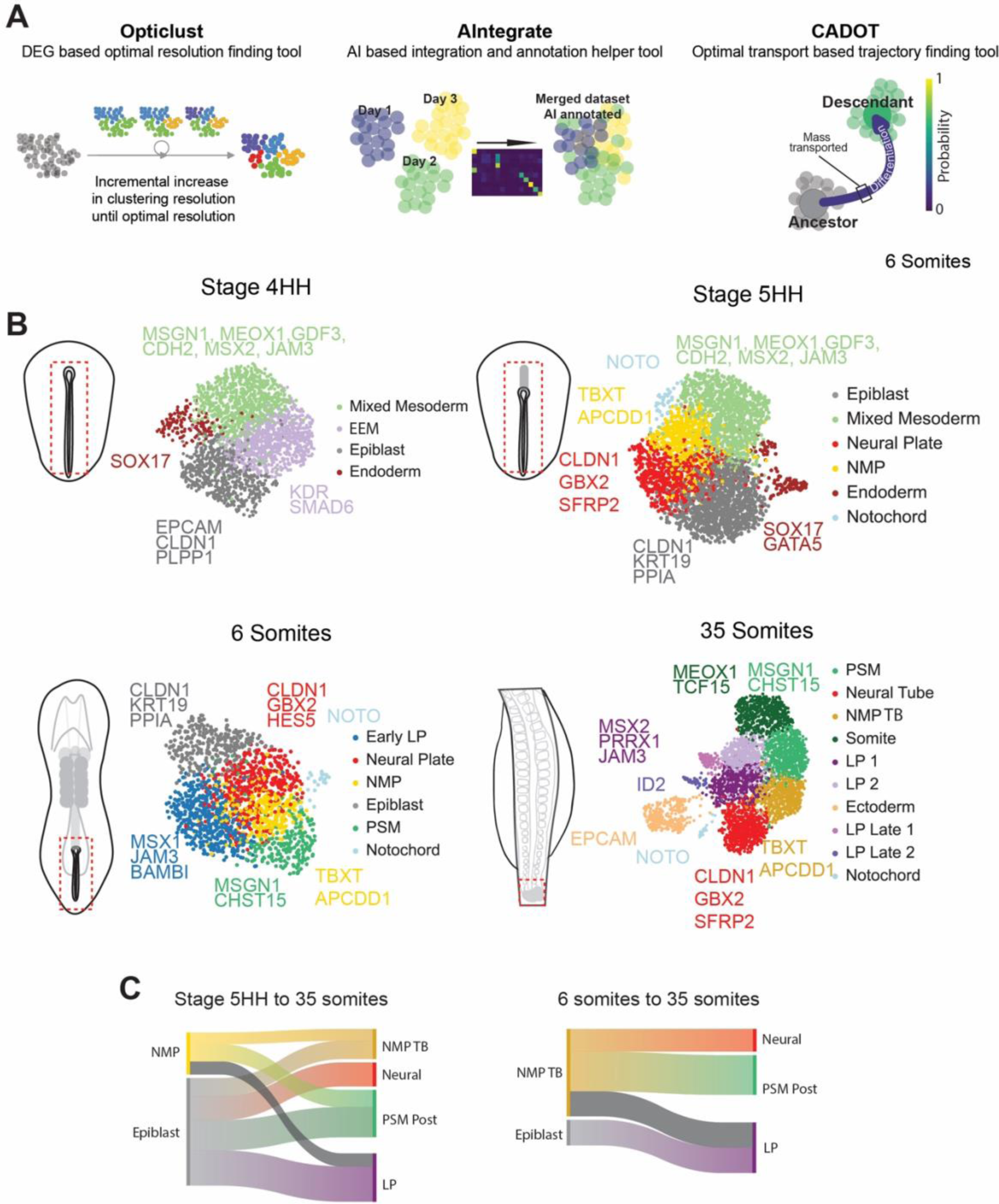
Developmental trajectory of the epiblast. (A) Trajectory analysis pipeline: Each dataset is annotated based on optimal clustering resolution (Opticlust), annotation of integrated datasets is based on a classifier (AIntegrate), and trajectories between every cluster of every dataset are identified using optimal transport (CADOT). (B) UMAP embedding of the single cell RNA sequencing datasets from the anterior PS region from HH4, HH5, 6 somites, and tail bud region from 35 somites chicken embryos after bkknn batch correction. Colors indicate clusters identified by Leiden and cell-type annotations using Opticlust. n=3 biological replicates for each stage. (C) Sankey diagram representing cell type transitions identified with CADOT. Transitions are computed between two developmental windows from stage 5HH to stage 35 somites (left) and stage 6 somites to stage 35 somites (right). The width of the transitions is proportional to the mass transported. Colors correspond to those of clusters shown in B, except for dark grey representing new lineage predictions.

We used Opticlust to investigate the cellular composition of the epiblast and the mesoderm in our dataset. Up to the 6-somite stage, the epiblast forms the superficial epithelial layer adjacent to the PS, from which mesodermal derivatives will ingress. At stage 4HH, the only other epithelial layer of the embryo is the endoderm that lines its ventral part. At this stage, we identified one cluster enriched in genes such as *SOX17* indicating that it corresponds to endodermal cells (Figure 1B, Supplementary Figure 2A). Another epithelial cluster, characterized by expression of *EPCAM, CLDN1, PLPP1* most likely represents the superficial epiblast (Figure 1B Supplementary Figure 2A). This cluster does not express the primitive streak marker *TBXT* but expresses *CDH1*, *POU5F3* (*OCT4*),^31^ and *SFRP2* (Supplementary Figure 2B). This combination of genes characterizes the epiblast cell state of the human Carnegie Stage 7 ^32^ which represents an approximately equivalent developmental stage to chicken stage 4HH. We thus named this cluster “epiblast”. This epiblast cluster was still present at stage 5HH (Figure 1B, Supplementary Figure 2C). At stage 5HH, we also detected another *CLDN1*-positive epithelial cluster which also expresses many genes enriched in NMPs (*TBXT, GJA1, DLL1, APCDD1, WLS, CDH2, ATP2B1, APLP2, WNT5A, WNT8A*) ^22^. This cluster, which we called NMP, likely corresponds to the anterior-most epiblast adjacent to the Node and anterior PS where the NMPs are first detected at this stage ^22^. The third epithelial cluster detected at stage 5HH corresponds to the endoderm (*SOX17* and *GATA5*). We also identified a fourth epithelial cluster expressing genes associated with Neural Plate identity (*GBX2, HES5*) (Figure 1B, Supplementary Figure 2C). At the 6-somite stage, the epiblast, NMP, and neural clusters were still present (Figure 1B, Supplementary Figure 2D). At the 35-somite stage, we could not detect any epiblast cluster except for an NMP cluster which we named NMP/TB since these cells localize in the tail bud at this stage (Figure 1B, Supplementary Figure 2E). A neural (*CLDN1, GBX2, SFRP2*) cluster was also identified at the 6 and 35-somite stages while a surface ectoderm (*EPCAM, KRT7, KRT19*) cluster was found at the 35-somite stage.

We also observed clusters exhibiting identities of ingressed mesodermal cells. At stage 4HH, opticlust identified an extraembryonic mesoderm (*GATA5, KDR, SMAD6*) and a mixed mesoderm cluster expressing both paraxial identity markers (*GDF3*, *MSGN1, MEOX1)* and lateral plate mesoderm identity markers *(JAM3, MSX2)* (Figure 1B, Supplementary Figure 2A) similar to the anterior somite cluster of mouse embryos at equivalent stages^33^. From stage 5HH on, Opticlust detected a Hensen’s Node/notochord (*CHRD, NOTO)* cluster. At the 6-somite stage, the LP and PSM identities segregated into 2 clusters (Figure 1B; Table 1, Supplementary Figure 2D). Opticlust identified a PSM cluster (*MSGN1, CHST15*) and an early LP cluster expressing the epithelial marker *KRT18* (Table 1) as well as the LP markers (*MSX1/2, JAM3, BAMBI*) probably corresponding to newly formed LP, which is epithelial at this stage. At the 35-somite stage, we identified mesodermal clusters including posterior PSM (*MSGN1*), anterior PSM/somite (*MEOX1, TCF15*), and LP (*MSX2, PRRX1, JAM3, ID2*).

To confirm the consistency of annotations amongst individual datasets and compare their similarity across developmental time, we developed a pipeline called AIntegrate (Figure 1A). This tool utilizes a machine learning classifier to identify similarities between clusters at different timepoints and then displays a projection of the clusters at each timepoint on the merged UMAP containing all developmental times (Figure 1A, Supplementary Figure 3A-B). It allowed us to confirm the identity of clusters found by Opticlust in the merged dataset containing all stages (4HH, 5HH, 6 somites, and 35 somites) (Supplementary Figure 3A). Our results suggest that from the onset of PS regression at stage 4HH, the region of the epiblast fated to give rise to non-axial mesoderm contains only two cellular states including an NMP and a non-NMP state. The non-NMP epiblast is transient and disappears after the 6-somite stage. Using a classifier trained on a mouse embryo atlas, we found good agreement between our cellular states and those of the atlas (Supplementary Figure 4A). We identify a mixed mesoderm cluster similar to the cluster of cells contributing to the anterior-most somites recently identified in mouse (Supplementary Figure 4A)^34^. These results suggest a mixed lateral/paraxial mesoderm identity of the first embryonic mesoderm cells produced after the beginning of PS regression. Thus, this clustering analysis suggests that the distinct mesodermal identities of epiblast territories are only acquired following ingression of multipotent precursors, probably in response to regional cues. Thus, the regionalized organization of the epiblast predicted from the fate maps (ie chordal, neuro-mesodermal, paraxial, intermediate, lateral, and extra-embryonic mesoderm) (Supplementary Figure 1) is not reflected at the transcriptome level in our analysis.

### Developmental plasticity of NMP and LP epiblast precursors in the chicken embryo

We next performed heterotopic and heterochronic transplantations of small regions of the epiblast of 1 to 3-somite chicken embryos to evaluate their developmental plasticity (Figure 2; Supplementary Figure 5). We transplanted a small domain from the anterior epiblast of the PS (NMP region), from Hamburger & Hamilton (HH) ^35^ stage 7-8 donors expressing GFP ubiquitously, into the prospective LP territory, at the mid-streak level of stage 5-6 HH unlabeled hosts (Graft1: G1) (Figure 2A). 20 hours after the graft, in 10 out of 10 embryos, the GFP-positive cells localized in the LP tissue indicating that grafted cells differentiated according to their novel environment (Figure 2B-E, Supplementary Figure 5A-B). We next assessed the identity of the transplanted GFP cells by immunofluorescence using known markers of the NMPs (SOX2^+^/TBXT^+^) ^36^, paraxial mesoderm (MSGN1^+^) ^22^ and LP (pSMAD1/5/9^+^) ^37^ ^37^ fates (Figure 2C-E). Cells of the NMP territory grafted in the LP domain produce LP-like descendants that are pSMAD1/5/9 positive (Figure 2C-D) but MSGN1 negative (Figure 2E). In contrast, isotopic and isochronic grafts of similar fragments gave rise to expected descendants including paraxial mesoderm and tail bud cells (Supplementary Figure 5D). Thus, epiblast cells of the NMP domain can change fate and give rise to LP cells when grafted into the LP progenitor region of the PS.

**Figure 2:**
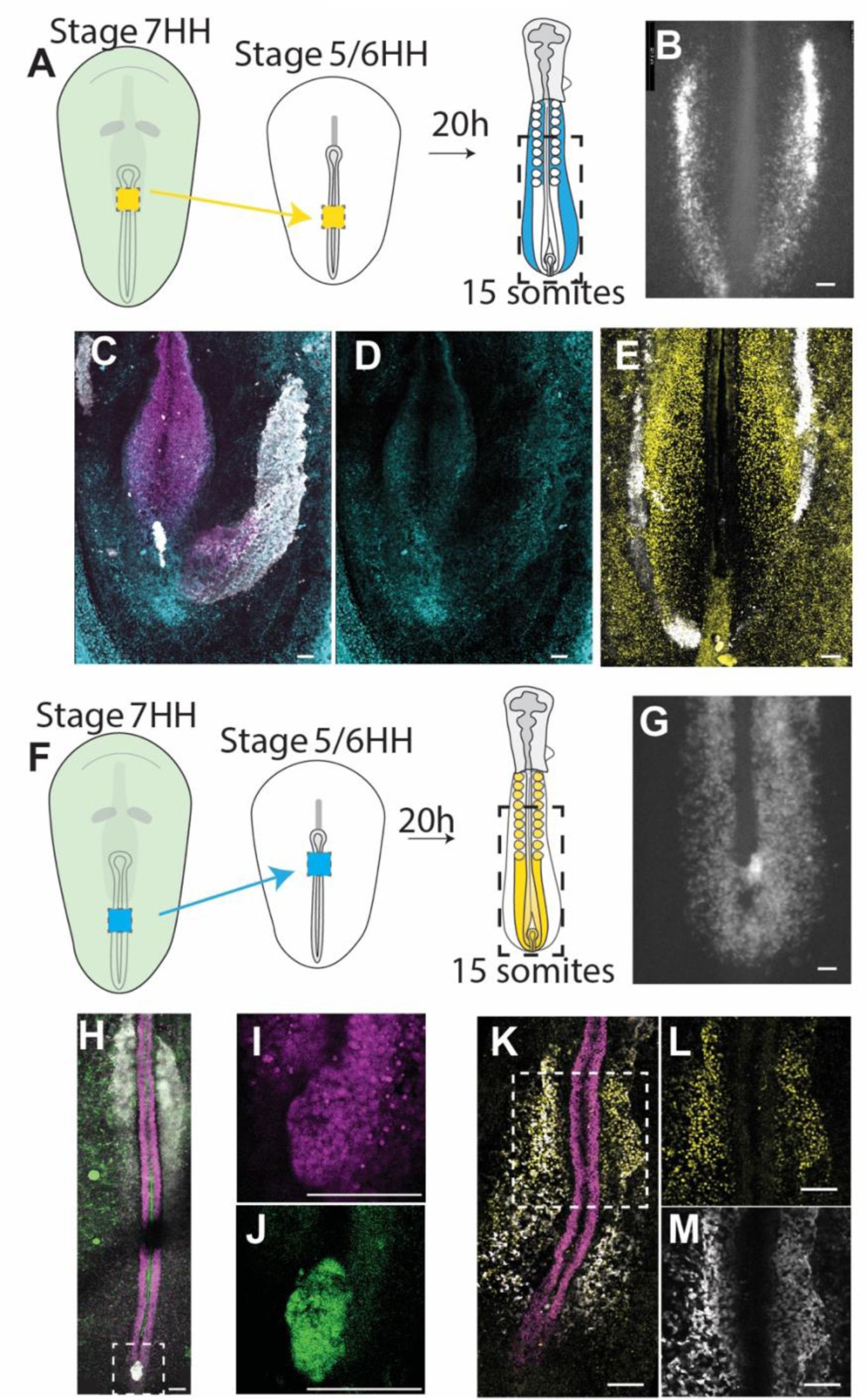
Developmental potency and transcriptional plasticity of epiblast cells. (A) Diagram showing type 1 grafts in which a fragment of the NMP progenitor domain (yellow) of stage 7-8 HH donor is transplanted into the LP progenitor domain of stage 5-6 HH host and the resulting localization of the GFP positive cells 20h after the graft (in blue, right). (B) Posterior region of an embryo transplanted as described in A incubated overnight, showing the localization of the transplanted cells which express GFP (boxed region in A). Bar=100µm (C and D) Confocal images of immunostainings on type 1 grafts 20 h after the graft showing the expression of the NMP/neural tube marker SOX2 (purple, C), and the localization of GFP cells (white, C), and the LP marker pSMAD1/5/9^+^ (blue, C-D) (E) Confocal image of immunostainings on type 1 grafts 20 h after the graft showing the expression of the paraxial mesoderm marker MSGN1 (yellow), and the localization of GFP cells (white). Bar= 100µm. (F) Diagram showing type 2 grafts in which a fragment of the LP progenitor domain (blue) of stage 7-8 HH donor is transplanted into the NMP progenitor domain of stage 5-6 HH host and the resulting localization of the GFP positive cells 20h after the graft (Yellow, right). (G) Posterior region of a embryo transplanted as described in F incubated overnight, showing the localization of the transplanted cells which express GFP (boxed region in F). Bar= 100µm (H) Confocal image of dorsal view of type 2 graft after a 20 h incubation showing the expression of SOX2 (purple), and the localization of GFP cells (white). Bar= 100µm. (I and J) Higher magnification of the boxed region in (H). SOX2 (purple), GFP (green). Bar= 100µm. (K) Confocal images taken from the ventral side of immunostainings on a type 2 graft 20 h after the graft showing the expression of SOX2 (purple, K), MSGN1 (Yellow, K-L), and the localization of GFP cells (white, K, M). Bar= 100µm. (L and M) Higher magnification of the boxed region in (K). Bar= 100µm.

We also performed the reverse type of grafts (Graft 2: G2) by transplanting a small fragment of the epiblast from the mid-streak level (corresponding to the LP progenitor domain) from a stage 7-8 HH GFP-expressing embryo to the anterior-most region of the PS (NMP domain) of a stage 5-6 HH host (Figure 2F, Supplementary Figure 5C). In control isotopic and isochronic grafts, these fragments give rise to LP descendants (Supplementary Figure 5D). Following a 20-hour reincubation period, in G2 grafts, GFP-positive cells integrated the paraxial mesoderm in 12 out of 12 grafted embryos (Figure 2G, Supplementary Figure 5B-C). In 10 out of 12 embryos, GFP cells were retained in the NMP domain of the tail bud (Figure 2H-J, Supplementary Figure 5C), suggesting that they can self-renew. The transplanted LP progenitors express NMP markers (SOX2^+^/TBXT^+^) (Figure 2H-J) and produce MSGN1-positive descendants when grafted into the NMP domain (Figure 2K-M). Few GFP cells were found in the neural tube in the G2 grafts. However, such was also the case in 5 out of 6 controls, where we performed isotopic and isochronic grafts of the same NMP territory from a stage 7-8 HH GFP donor into a stage-matched unlabeled host (Supplementary Figure 5E). This is most likely due to the short reincubation period using *ex ovo* culture as we previously showed that the bipotentiality of NMPs is mostly expressed at later developmental stages ^22^. Cells of the LP progenitor region appear therefore to switch to an NMP fate when grafted in the anterior epiblast. Together, these experiments show that cells from the NMP and LP domains of the PS epiblast are not committed to a specific cell fate at these stages. They respond to their novel environment by losing their original identity, acquiring markers appropriate to the fate of their novel location.

### Bioinformatics analysis reveals unexpected developmental trajectories of the epiblast/NMP cells

To understand how mesodermal cell fates diversify from the epiblast/NMP state, we developed a novel bioinformatics approach to study their developmental trajectories. We examined the relationship between cellular states identified in our scRNAseq dataset at all timepoints. To do so, we developed a program using the optimal transport model (Waddington-OT ^38^) to predict cluster-to-cluster transitions. We call this trajectory hypothesis generator program CADOT (Cluster ADjusted Optimal Transport) (Figure 1A). Cells from all timepoints are distributed based on their transcriptional similarities in a 2D environment (ie UMAP), and all the calculated transitions between clusters are displayed on the graph using arrows that go from one cluster (ancestor) to another (descendant). Additional information can be studied including the transition probability and the quantity of cells transported from one cluster to another. We chose stringent filter parameters such as unlikely transitions like anterior mesoderm/neural lineage transition are not represented. Visualization of the trajectories can be achieved using a Sankey diagram where the width represents the quantity of mass transported during a transition (Figure 1C) or directly in the UMAP (Supplementary Figure 3C) with arrows indicating both mass transport (width of the arrow) and probability of the transition (color).

Using these filtering parameters, we first computed the hypothetic ancestors/descendants’ relationships for epiblast cell states and their putative descendants by sub-clustering the ancestor cells: epiblast, NMP, and NMP/TB and descendant cells: neural, mixed mesoderm, LP, PSM Post, EE of stages 5HH, 6 somites, and 35 somites (Supplementary Figure 3A-B). We analyzed lineage transitions from the epiblast to mesodermal clusters by computing transitions across two developmental windows, from stage 5HH to 35 somites, and from 6 somites to 35 somites, to infer the immediate and long-term lineage trajectories of epiblast cells. This predicted well-known transitions associated to the multipotency of epiblast and NMP cells identified in classical avian and mouse lineage studies. For instance, interrogating CADOT to predict lineage trajectories from stage 5HH to 35 somites (Figure 1C, Supplementary Figure 3C) reveals that the epiblast cluster gives rise to PSM post, LP clusters, NMP/TB and neural clusters. In addition, CADOT predicts a departing node from the NMP cluster to the NMP/TB cluster, which describes the maturation of NMP cells ^22^. CADOT also identifies a transition from the NMP to the PSM fate recapitulating the short-term lineage fate mapping of this region in avian and mouse embryos ^15,22^. From 6 somites to 35 somites (Figure 1C, Supplementary Figure 3C), CADOT also recapitulates known lineage transitions happening at these developmental times. From the NMP/TB node, which is the largest departing node, CADOT identifies transitions toward the PSM and neural clusters recapitulating the expected neuro-mesodermal bipotency of NMPs. We also detected the expected transition from epiblast to LP. Altogether, this shows that CADOT accurately predicts expected developmental trajectories from the epiblast.

In parallel to these known transitions, CADOT also predicts previously undescribed trajectories departing from the NMP and the NMP/TB clusters with high probability. Specifically, we observed transitions from the NMP or NMP/TB to the LP fate (Figure 1C). This represents a truly novel differentiation trajectory for NMPs. Additionally, from stage 5HH to 35 somites (Figure 1C, Supplementary Figure 3C), lineage trajectories depart from the non-NMP epiblast to neural and NMP/TB, suggesting that this epiblast can also give rise to these two lineages. These hypothetic trajectories are consistent with the developmental potency of epiblast observed in our grafts. Thus, our results hint at considerable plasticity of cells of the anterior and posterior epiblast domains. Overall, our advanced trajectory pipeline combining Opticlust, AIntegrate, and CADOT, predicts known and novel ancestors/descendants’ lineage trajectories. The most surprising results from CADOT predictions is the suggestion that both NMP and non-NMP epiblast cells can give rise to more cell fates than the neural and paraxial mesoderm lineages.

### Single-cell lineage tracing identifies multipotent progenitors in the posterior epiblast domain

To validate CADOT predictions, we investigated the lineage potency of single cells of the LP progenitor domain of the epiblast which lies at the mid-streak level between stages 4HH to 8HH. To track the fate of single cells of this region, we used a lineage tracing strategy based on the Brainbow-derived MAGIC markers ^39^. We performed local co-electroporation of plasmids expressing a self-excising Cre recombinase and the Nucbow transgene together with the TolII transposase to drive transgene integration in epiblast cells. This strategy allows to permanently mark cell nuclei with a specific color code generated by the unique combination of different fluorescent proteins triggered by random recombination of the Nucbow cassette. This color code is then stably transmitted to each daughter cell and can be retrieved by confocal imaging and quantification of the color hues of cells in the electroporated embryos after a 48h reincubation period. Labeling cells at the mid-streak level at stage 5HH gave rise to descendant cells in the mesodermal, neural, and LP tissues (Figure 3A-D, Supplementary Figure 6). Most of the cells were labeled in the region posterior to the forelimb at all anteroposterior (AP) levels of the thoracic, lumbar and sacral regions. (Figure 3B, C Supplementary Figure 6) with many cells sharing the same color code in the paraxial mesoderm and LP (Figure 3B-E). Also, we found cells labelled in the neural tube (Supplementary Figure 6). These cells present color codes which could be distributed across several tissues such as neural tube and somite, or neural and lateral plate. In 20 % of the 31 clones analyzed, we found cells with the same clonal identity in the neural tube, paraxial mesoderm and LP (Figure 3E). Thus, single cells of the 50% level of the PS at stage 5HH, which are classically considered fated to give rise to LP can give rise to descendants in multiple tissues of the posterior embryo including the neural tube. In contrast, labeling the epiblast of the mid-streak level at stage 7-8 HH generated mostly LP cells (Figure 3F-G). Virtually no labeled cells were found in the neural tube or paraxial mesoderm when electroporation was performed at this stage. Thus, at stage 7-8 HH, cells of the mid-streak level are specified to the LP fate. These cells may however not be committed to this fate yet as they retain the potency to give rise to multiple germ layers as predicted by CADOT (Figure 1C) and shown with the type 2 grafts (Figure 2E). Overall, our single-cell lineage tracing analysis indicates that at stage 5HH, epiblast cells of the mid-streak level are multipotent, generating neural, paraxial mesoderm, and LP lineages in the posterior embryo. In contrast, epiblast cells of the same PS level at stage 8HH mostly generate LP descendants.

**Figure 3:**
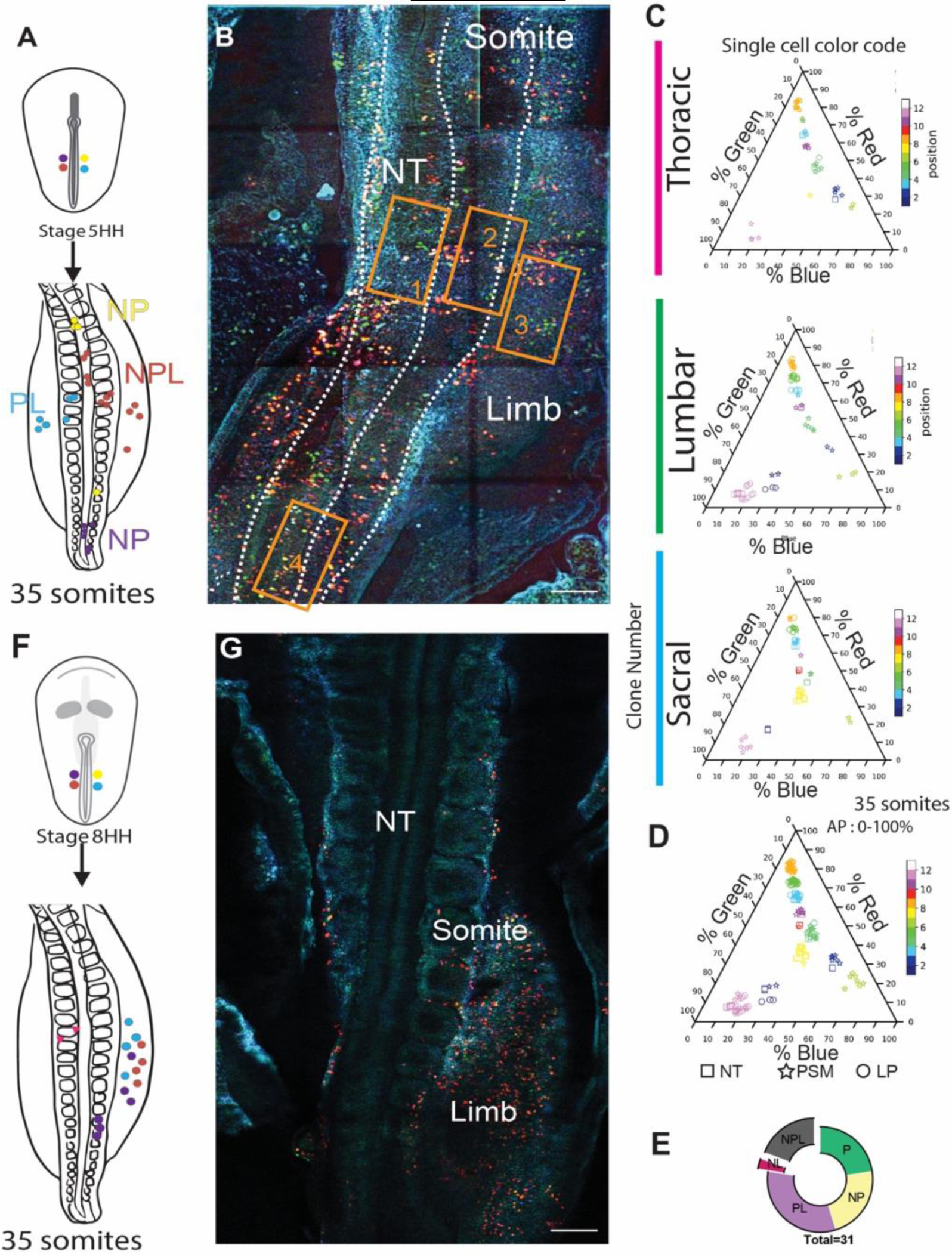
Single-cell lineage tracing identifies multipotent progenitors in the posterior epiblast. (A) Diagram showing the electroporated region in the presumptive LP progenitor domain of the epiblast at stage 5HH (top) and the stage at which embryos were harvested for analysis (bottom)(n = 7). Color dots show example of single cells with a specific color code and their localization at electroporation and at harvesting time. (B) Confocal z-projection of a stage 17HH embryo acquired using three separated laser paths to retrieve the Nucbow color codes genetically encoded as described in Loulier et al., 2014. Stippled lines delimit the neural tube and somite regions. NT: neural tube. Numbered orange boxes correspond to higher magnifications shown in Supplementary Figure 6. Bar = 100µm. (C and D) Triplot diagrams showing the distribution of descendants of cells labeled with different Nucbow combinations along the thoracic, lumbar and sacral AP axis (C) or merged (D) of eleven clones in a representative stage 17HH embryo. Each symbol represents a cell identified based on the percentage of red, blue, and green expressed. The symbols are colored based on their clonal identity. Squares: neural cells; stars: paraxial mesoderm cells; circle: lateral plate cells. (E) Quantification of clones frequency: Paraxial mesodermal (P, green) Paraxial and Lateral plate mesoderm (PL, purple), bipotent Neural and Paraxial mesoderm (NP, gold), bipotent Neural and Lateral plate (NL, red) and tripotent Neural, Paraxial mesoderm and Lateral plate (NPL, grey) at stage 17HH (n = 31 clones, in seven embryos). (F) Diagram showing the electroporated region of the presumptive LP progenitor domain of the epiblast at stage 8HH (top) and the stage at which embryos were harvested for analysis (bottom) (n = 5). Color dots show single cells with a specific color code and their localization at electroporation and at the harvesting time. (G) Confocal z-projection of a stage 17HH embryo acquired using three separated laser paths to retrieve the color codes genetically encoded. Bar = 100µm.

### Analysis of tissue and cell dynamics identifies boundaries between territories of the epiblast

Our lineage tracing analysis indicates that mid-streak epiblast cells at stage 5HH can give rise to lineages normally derived from more anterior levels such as neural tube or paraxial mesoderm. One possible explanation for this surprising observation is that epiblast cells can move within the plane of the tissue to relocate more anteriorly during PS regression. To investigate cell movement and tissue remodeling in the epiblast, we used the dynamic morphoskeleton analysis method^40^. This strategy allows to project onto the embryo the future dynamic behavior of cells. We first generated movies of the PS region of transgenic chicken embryos expressing GFP ubiquitously from stage 4^+^HH to stage 8HH (Supplementary Figure 7A-C). We then quantified tissue velocity fields using Particle Image Velocimetry (PIV) analysis to identify regions where cells which are initially closely located diverge with time (repellers, Forward projection, FW) or where initially distant cells converge (attractors, Backward projection, BW) (Supplementary Figure 7 A-B). This analysis allows visualizing future cell dynamics on the initial tissue configuration in the embryo. We identified two significant repellers at stage 4^+^HH: one which delimitates the mesodermal territory of the epiblast from the neural plate anteriorly and one that marks the boundary with the non-neural ectoderm laterally (Supplementary Figure 7A, D, F). We also identified one main attractor corresponding to the PS/ingressing zone towards which epiblastic cells converge (Supplementary Figure 7A, E-F). Thus, our morphoskeleton analysis correctly delimitates the epiblastic territory of mesodermal precursors from future ectodermal domains. This analysis did not reveal any repellers separating the future mesoderm territories of the epiblast along the AP axis.

To increase the resolution of our analysis, we generated 10h movies of embryos labeled with the vital dye nuclear red (Figure 4A). This allows to sparsely label the nuclei of epiblast mesodermal progenitors to identify individual cell tracks. Using these tracks with the morphoskeleton pipeline, we could identify 2 stable repellers forming orthogonally to the PS (Figure 4A-E). We named these repellers Boundary 1 and 2 (B1, B2) based on their timing of appearance during embryonic development. B1 appeared at mid-streak level, 200 minutes after the beginning of PS regression (Figure 4 F-H, N-O). The location and time (stage 6HH) at which this boundary forms maps to the level of the expected segregation of the embryonic and extraembryonic mesodermal domains of the epiblast. Thus prior to B1 formation, epiblast cells at the mid-streak level can freely move in this region (Figure 4G-I). The presence of the B1 repeller, indicates that after stage 6HH, cells remain on each side of B1, becoming confined to the extraembryonic and embryonic (LP) mesoderm domains of the epiblast. B2 forms 312 minutes after the beginning of PS regression (stage 7-8 HH) at the 70% PS level (Figure 4J-O). The location and time at which this boundary forms suggest that B2 separates LP from PM progenitors in the epiblast. After B2 forms around stage 7-8 HH, cells remain confined on either side of the repeller and will give rise either to LP or PM descendants. Importantly, this dynamics of B2 formation parallels the fate restriction observed in our lineage tracing analysis of mid-streak progenitors. Thus, while cells of the epiblast remain uncommitted as seen by our scRNAseq analysis and graft experiments, our results suggest that their physical confinement to a specific epiblast domain controls their specification to a specific lineage. This morphoskeleton analysis provides a dynamical view of the fate map formation in the epiblast complementing the static fate maps established by classical lineage tracing methods.

**Figure 4:**
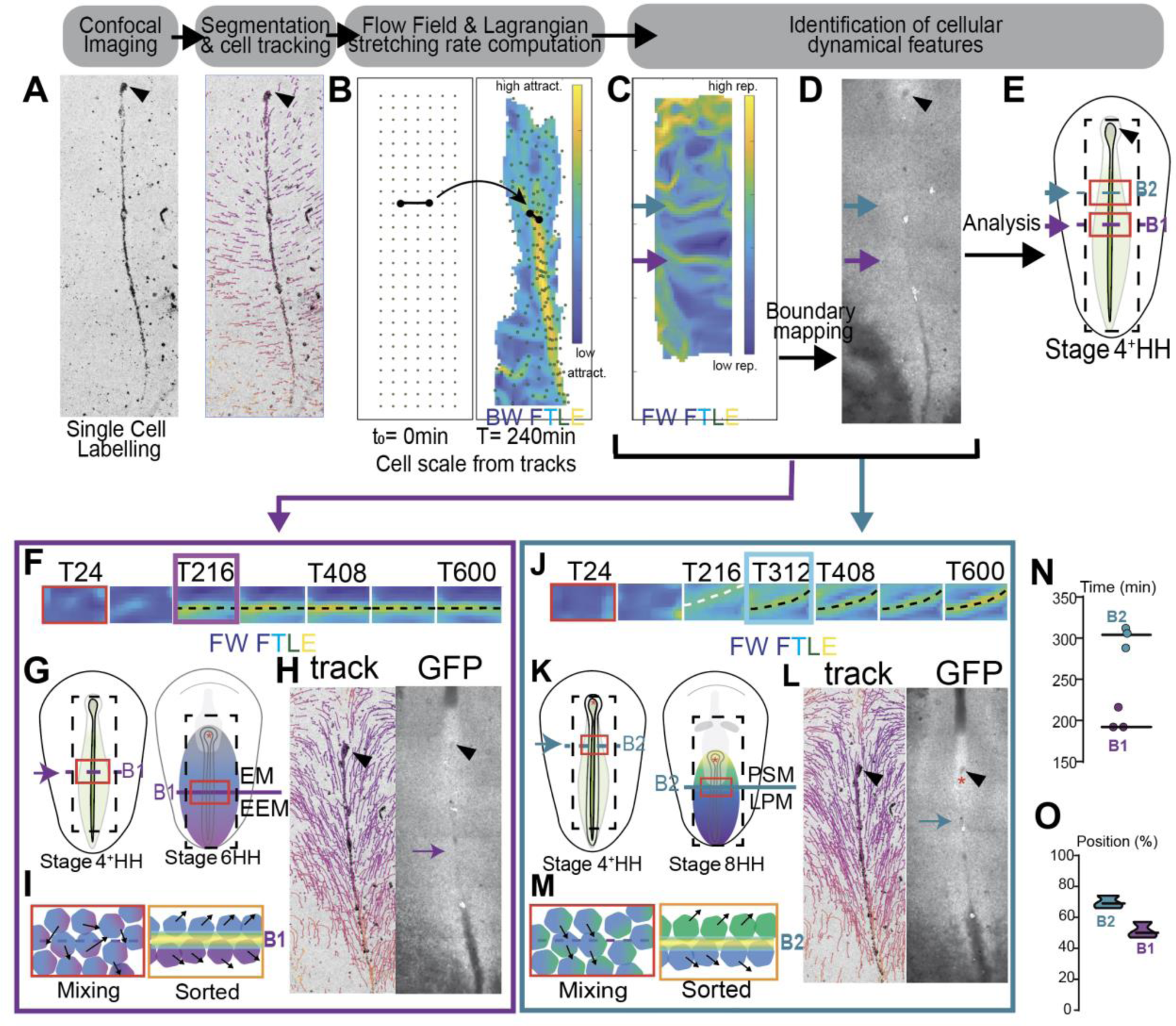
Dynamic Morphoskeleton analysis of epiblast cell dynamics identifies boundaries between precursor territories. (A to E) Workflow to identify repellers and attractors using the dynamic morphoskeleton on cell tracks. Representative images showing the raw confocal image of nuclearRed labeled stage 4HH embryos (A) analyzed with the dynamic morphoskeleton (B-C). (B) Grid deformation (green dots) and the Backward FTLE field whose high values (yellow) locate the attractor. (C) Forward FTLE field whose high values (yellow) locate repellers. (D) Projection of the identified repellers on a stage 4HH embryo (E) Scheme representing the position of the B1 and B2 boundaries identified as repellers by the morphoskeleton pipeline. (F) Close-up of the FW FTLE color-coded image showing the dynamic formation of the Boundary 1 (black dotted line) over time. (G and H) Representative diagram showing the localization in space (G), the cell trajectories and the raw image (H) at the time (purple square in F) of boundary formation. (I) Diagrams showing cell shuffling before boundary formation and cell sorting after the boundary appears, forming two distinct tissue compartments on each side. (J) Close-up of the FW FTLE color-coded image showing the dynamic formation of the Boundary 2 (black dotted line) over time. The white dotted line in J shows an unstable repeller. (K and L) Representative diagram showing the localization in space (K), the cell trajectories and the raw image (L) at the time (blue square in J) of boundary formation. (M) Diagram showing cell shuffling before boundary formation and cell sorting after the boundary appears, forming two distinct tissue compartments on each side. (N and O), Quantification of the time (N) and location (O) of formation of B1 and B2 boundaries. n=3 embryos. Data are represented as mean ± SEM.

### Displacement of epiblast progenitors depends on cell intercalation following cell division

Our morphoskeleton analysis did not identify repellers in the epiblast before stage 6HH when cells can contribute to several distinct lineages. This is consistent with the idea that cells can freely move in the tissue at these stages. However, since the epiblast is a pseudostratified epithelium, its cells are expected to be constrained in their movements. One possibility is that cell division combined or followed by cell intercalation allows the displacement of daughter cells within the epiblast. This could allow them to relocate and ingress in different domains, thus giving rise to different fates. To test this hypothesis, we performed live imaging of the epiblast of transgenic quail embryos expressing a nuclear red (H2B-mCherry) and green membrane fluorescent reporters (membrane-GFP)^41^. We quantified the number of cell division events from 0 to 2 hours, 2 to 4 hours and 4 to 6 hours after the start of PS regression. We found that the number of cell division events progressively decreases during the first hours of PS regression (Figure 5A). Measuring the distance between sister cells at the end of cell division identified two distinct cell division patterns: non-intercalating sister cells (ie: 0 to 15 µm between daughter cells) and intercalating sister cells (more than 15 µm between daughter cells) (Figure 5B-D). Intercalation events upon cell division occurred mostly during the first 4 hours of PS regression (between stage 5 to 7-8 HH) in the first 700 µm of the anterior PS region (62 to 100% PS, Figure 5D, left). We observed limited number of intercalating daughter cells upon cell division in the more posterior region of the epiblast (750 to 1450 µm below the node, ie 20 to 58% PS, Figure 5D, right). These results support the existence of different cell division patterns in the epiblast during the early stages of PS regression.

**Figure 5:**
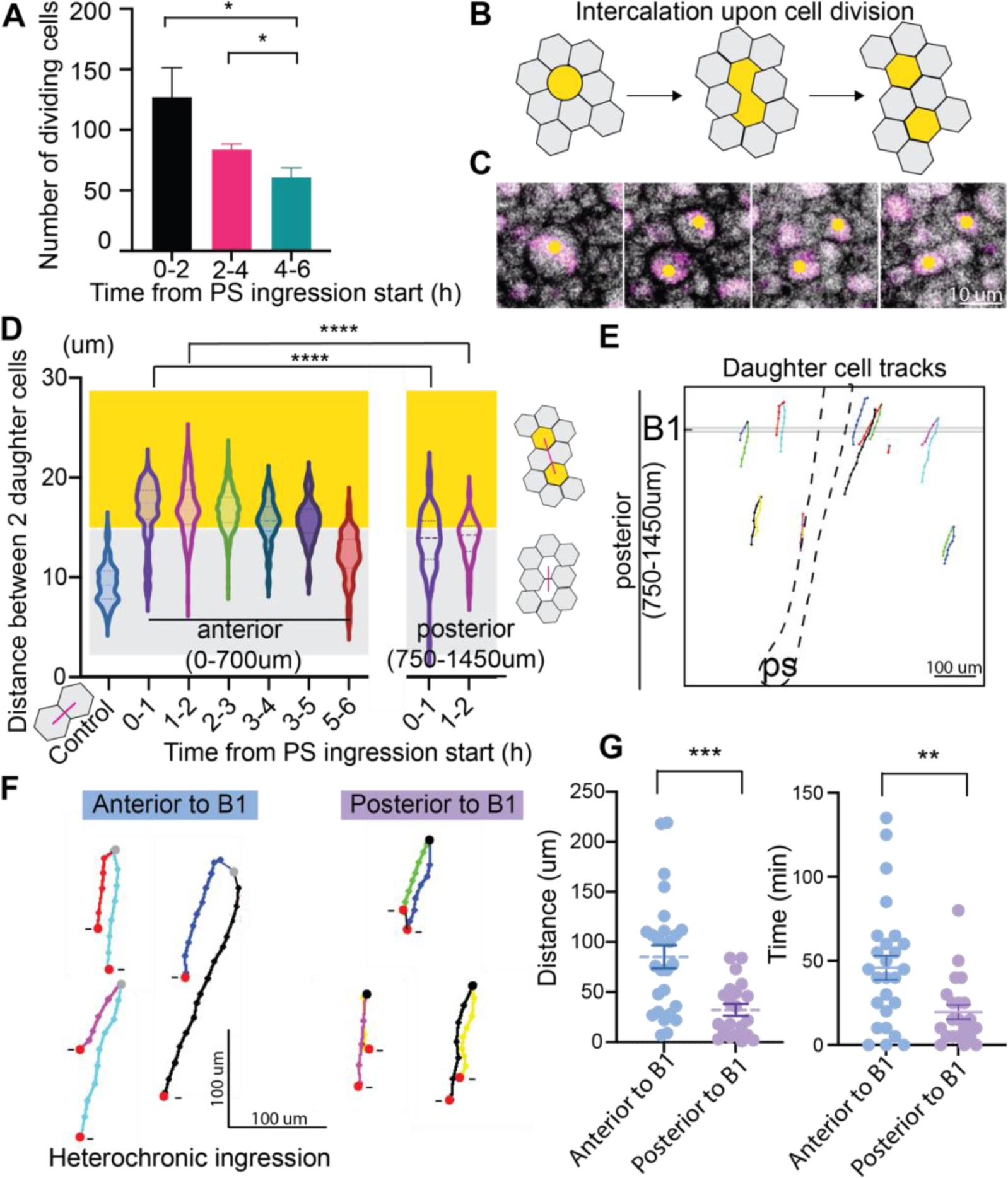
Displacement of epiblast progenitors is driven by cell intercalation following cell division. (A) Number of dividing cells in the epiblast from the beginning of PS regression (0 hour, 2h binning window). (N=2 embryos, n=1109 division events) Data are represented as mean ± SEM. (B) Diagram showing cell intercalation upon cell division (yellow cell) in the epiblast (grey). (C) Representative image of intercalation upon cell division using transgenic quail embryos expressing nuclear red and membrane GFP. Scale bar = 10 μm. (D) Quantification of the distance between daughter cells after division in the anterior and posterior PS over time. 0 hours marks the start of PS regression. Distance in the control represents the distance between 2 adjacent cells in the epiblast. Yellow zone shows where measurements are superior to twice the mean distance between cells in the epiblast (ie: intercalation of 1 cell). N=2 embryos, n=1149 division events. Data are represented as mean ± SEM. (E) Representative manual tracking of daughter cells dividing in the posterior PS region (750 to 1450 μm) where B1 forms (grey). (F) Single manual tracks of daughter cells after they divide anterior to the B1 location (grey dots, left) or posterior to the boundary region (black dots, right). Red dots show the end of sister cells tracking due to ingression (-). (G) Quantification of the distance (left) and time (right) between sister cells ingression coordinates for sister cells born anterior (blue) or posterior to B1 location (violet). N=2 embryos, n=47 cells. Scale bars = 100 μm. Data are represented as mean ± SEM.

To test if the displacement of daughter cells within the epiblast supports their relocation and ingression in different AP domains, we tracked the position of sister cells originating from a mother cell anterior (grey dots) or posterior (black dots) to the midstreak B1 location (grey line) until their ingression (0 to 240 min) (Figure 5 E-F). Cell tracks of sister cells emerging anterior to B1 ended up ingressing far away from each other while tracks from sister cells emerging posterior to B1 location ingressed closer to each other (Figure 5F). We quantified the distance and time difference between ingression coordinates of sister cells born anterior or posterior to B1 (Figure 5G). We find that cells born in the posterior streak region (posterior to B1), ingress on average 32 µm apart with a 19 min delay (Figure 5G). Anterior to the B1 level, we find a bimodal distribution of distance and time of ingression. Specifically, we observe that 30% of the sister cells ingress with a similar spatiotemporal dynamic as cells of the posterior PS region (16/52 cells ingress 50 µm apart within the first 30 minutes after they were born) (Figure 5G). In contrast, most sister cells born anterior to the midstreak level ingress from 80 to 218 µm apart with a delay between 50 to 135 min. Thus, sister cells born at or anterior to the midstreak level can end up in different presumptive territories given this ingression dynamics. Such sister cells can thus be exposed to distinct spatial and temporal cues that may explain their acquisition of different fates.

Thus, our results show that intercalation after cell division can affect the distribution of daughter cells within the epiblast during early stages of PS regression. Such displacements of progenitors in the epiblast may explain the multiplicity of fates of mid-streak epiblast cells observed in our cell lineage analysis.

### Anterior and posterior epiblast cells are exposed to distinct mechanical constraints

Our data indicate that despite being specified to the LP fate, cells of the mid-streak epiblast at stage 7-8 HH retain plasticity and can contribute to PM when grafted at a more anterior level. This therefore suggests that external factors control the fate of these progenitors. One such set of factors which has been poorly investigated is the mechanical environment of the epiblast at these stages. To investigate the role of mechanical constraints in cell fate determination, we first mapped tissue deformation using PIV analysis of the epiblast of stage 5HH to 9HH transgenic GFP embryos (Figure 6A). To do so, we divided the tissue into a grid of 80 µm squares and followed their deformation over time. In the anterior PS domain, we observed a change in grid shape from a square to an enlarged and elongated diamond shape without losing grid squares (Figure 6A). This indicated important anisotropic deformations without tissue loss consistent with the cell intercalation observed. This also suggests the existence of pulling forces from neighboring tissues in both the AP and ML directions (Figure 6A-B). The increased diagonal length of the diamond in this region of the grid indicates a greater deformation along the AP axis, marking the direction of anisotropy. Tissue loss seen by the disappearance of grid squares (pink dots) is visible anteriorly starting at 4 hours (T=264 min) after the start of PS regression. Thus, we find massive anisotropic tissue rearrangement within the peri-node region and the anterior PS during the early stages of its regression (Figure 6A-B). In contrast, limited anisotropic deformations were observed in the more posterior PS region indicating that the epiblast undergoes deformation without shearing at this location. As expected from the known ingression flow of posterior epiblast, grid squares flanking the posterior part of the PS start disappearing from the beginning of PS regression indicating progressive tissue loss (pink dots). Hence, anterior and posterior PS domains are under different mechanical constraints with an expansion/remodeling zone in the anterior epiblast around the node and an ingression zone in the posterior epiblast (Figure 6A-B).

**Figure 6:**
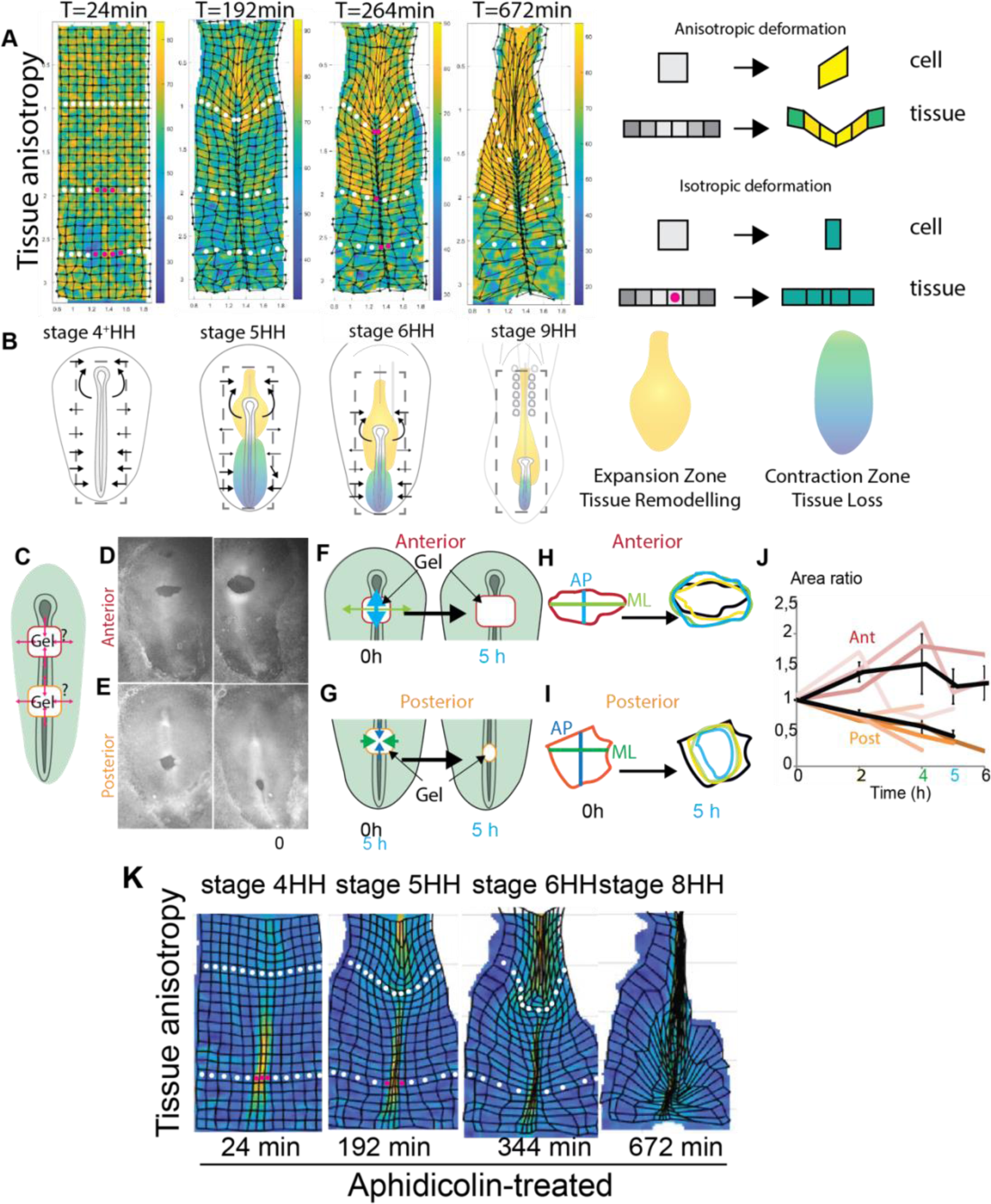
Anterior and posterior epiblast cells are exposed to distinct mechanical constraints. (A and B) Deformation maps (A) and corresponding diagrams (B) representing the region analyzed at each developmental stages showing the percentage of anisotropic tissue deformation. Data obtained from live imaging of transgenic chicken embryos expressing GFP at increasing time intervals (T) (grid size = 80μm) (n=3). Yellow domains in A label regions characterized by high anisotropic deformations ^40^. White dots show tissue remodeling zones and pink dots show tissue loss regions. (C to G) Diagrams (C, F, G) and representative images (D, E)) of the gel implants shapes at t=0h and t=5h after implantation. (H and I) Shapes delimitating the gel implant at t=0h (left) and overlay of the shapes over 5 hours in the anterior (H) and posterior (I) epiblast regions. The color coding shows the time after gel implantation 0h, black, 2h yellow, 4h green, 5h blue. (J) Quantification of the evolution of the area normalized to initial gel size at t=0h and over 6 h. Black curve is the mean curve of 8 embryos. Data are represented as mean ± SEM. (K) Representative images displaying the percentage of anisotropic tissue deformation (high values in yellow) at increasing time intervals to show tissue deformations in aphidicolin-treated transgenic GFP chicken embryos (grid size = 80μm) (n=3) White dots show tissue remodeling zones and pink dots show tissue loss regions.

We next directly probed the forces within the anterior and posterior epiblast by monitoring the deformation of soft alginate gels inserted into these two epiblast regions *in ovo* (Figure 6C, Supplementary Figure 8A-C). We first performed a small ablation of a squared fragment of tissue (without removing the endodermal layer) at different AP positions along the PS in GFP transgenic embryos. We next implanted a small piece of alginate gel in the region of the ablation and measured the shape changes of the implant (Figure 6C-J, Supplementary Figure 8). In the anterior epiblast, we observed significant tissue relaxation in the ML direction when cutting the tissue to place the gel implant indicating that this region is under high tension (Figure 6D, F, H, J). By quantifying the evolution of the shape of the gel implant over time, we measured a 50% increase in area 4 hours after placing the implant (Figure 6H-J). We observed an increase in length along the AP and ML axis with a bias toward a greater deformation along the AP axis (Supplementary Figure 8B-C). These deformations are consistent with pulling forces from neighboring tissues resulting in the anisotropic deformations described above (Figure 6A-B). When the implant was placed in the posterior epiblast domain, we observed a 50% decrease in the ML dimension after 5 hours indicating pushing forces from the ML tissues (Supplementary Figure 6C). In contrast, the gel AP length remained approximately constant (Figure 6E, G, I-J; supplementary Fig 6A-B).

We next used aphidicolin to test the role of cell division on the dynamics of tissue remodeling, (Figure 6K, supplementary Fig 9). Using PIV analysis on treated embryos, we found that this treatment decreases the percentage of local anisotropic deformation within the epiblast particularly around the node region compared to WT (Figure 6K). Thus, hampering cell division specifically impairs tissue remodeling in the anterior domain (around the node), but not posteriorly. Importantly, the morphoskeleton analysis on PIV measurement of aphidicolin-treated embryos identified similar attractor and repellor regions (supplementary Fig 9) suggesting that cell division inhibition does not impact mesoderm ingression. Thus, at the onset of PS regression, the anterior and posterior regions of the epiblast of the PS are exposed to different mechanical constraints. Pulling forces lead to epiblast remodeling anteriorly while ML compression associated with tissue ingression in the PS is observed posteriorly. Furthermore, cell division plays an important role in remodeling the anterior epiblast.

### Interfering with epiblast mechanical properties affects mesoderm progenitors’ fate

To directly test the impact of mechanical constraints on mesoderm progenitor fate decisions *in ovo*, we placed a filter paper on top of the right epiblast parallel to the PS of stage 7-8 HH embryos, and reincubated them for 5 to 12 hours (Figure 7A-B). We next tested how the filter affects tissue tension by cutting embryos along the PS in controls and in embryos implanted with the filter. We tracked the dynamics of the cut opening using a kymograph visualization (black stippled line, Figure 7C-D). In controls, the cut opening increased, consistent with the existence of pulling forces from the neighboring tissues in the anterior epiblast (Figure 7C). On the contrary, in embryos with a filter, limited relaxation is observed at the cut site (Figure 7D). Thus, placing the filter changes the mechanical properties of the epiblast, decreasing the ML pulling forces in the anterior region.

**Figure 7:**
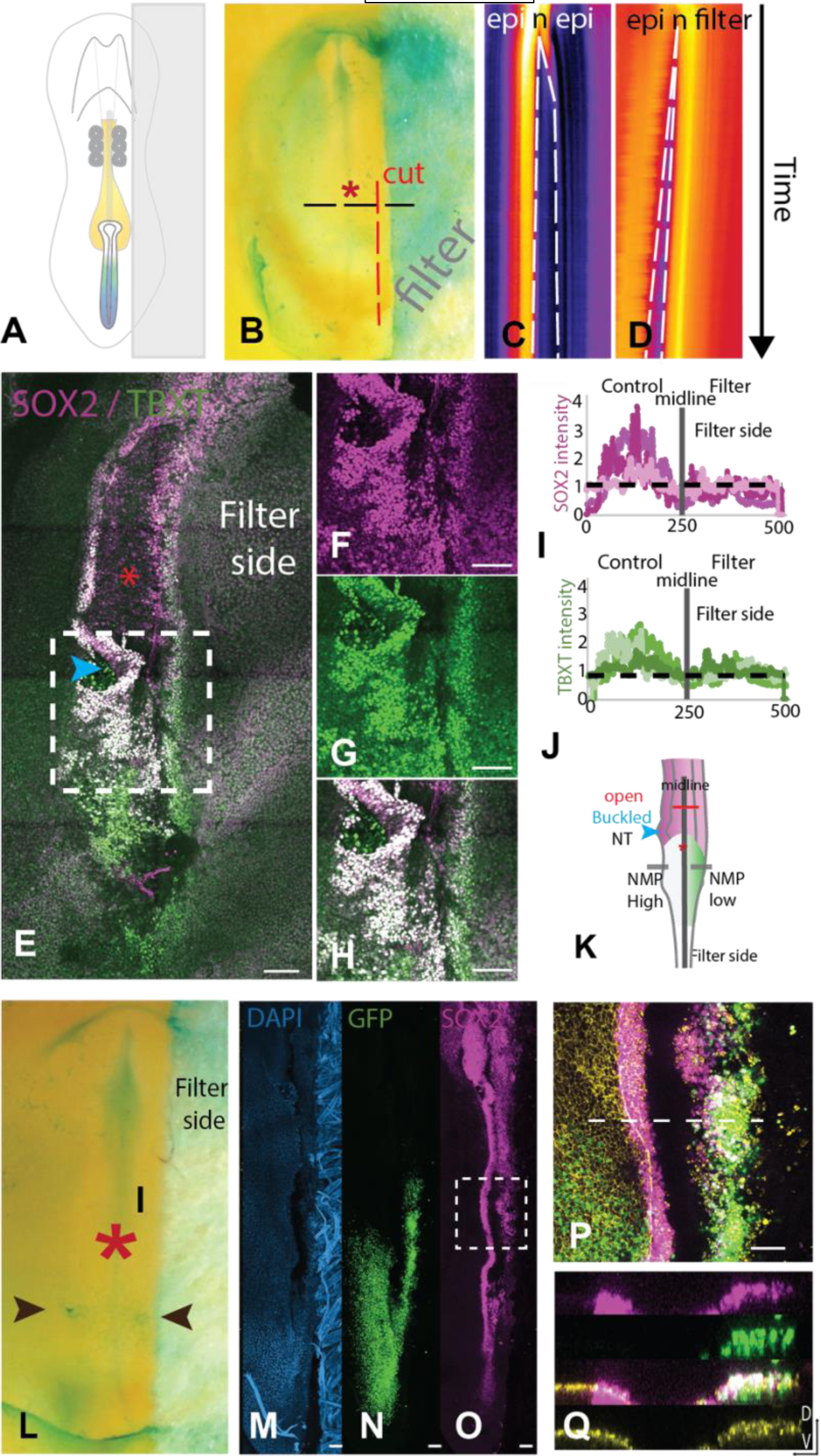
Interfering with epiblast mechanical properties alters the fate of mesoderm progenitors. (A) Diagram showing filter positioning to interfere with tissue mechanical properties in a stage 8HH embryo. (B) Stage 8HH chicken embryo with the filter in place. The red stippled line shows the location of the cut to test tension changes. The black stippled line shows the position of kymograph tissue analysis. (C and D) Kymographs showing relaxation after cutting the embryo in WT (left) and embryos with filter (right). (E) Representative image of an embryo reincubated overnight after filter implantation as shown in A. Immunochemistry against TBXT (green) and SOX2 (purple). Bar= 100µm (F and H) Detail of the region shown in the squared box in (E) labeled with antibodies against SOX2 (F), TBXT (G), and merge (H). Bar= 100µm. (I and J) Quantification of the fluorescence intensity of SOX2 (purple, I) and TBXT (green, J) across the NMP domain (dotted grey line in K) (K) Summary diagram of the effect of filter implantation on the NMP cell population. (L) Representative image showing the electroporation sites in the LP domain of the epiblast (arrowheads). (M to O) Representative images showing DAPI (blue) (M) indicating the position of the filter, the electroporated cells (green) (N), and SOX2 (purple) (O) in stage 8HH electroporated embryo incubated overnight after placing the filter on the right side. Bar= 100µm (P) Higher magnification of the neural tube region boxed in (O) showing SOX2 (purple), GFP (green) and F-actin (Yellow) staining. White indicates cells expressing both SOX2 and GFP. Bar= 100µm (Q) Transverse confocal sections at the level of hatched line shown in P. Same color code as in (P).

We next used this filter method (without cutting the embryo) followed by immunolabelling with TBXT and SOX2 antibodies to assess the identity of the descendants of the anterior epiblast cells exposed to this new mechanical environment. Placing the filter at stage 7-8 HH resulted in opened and buckled neural tube on the opposite side of the filter, indicating increased tension in the left side of the embryo (Figure 7E). Surprisingly, we observed an increased number of NMP cells on the side lacking the filter (Figure 7E-H). Analyzing the intensity of TBXT/SOX2, in the NMP domain shows high SOX2 expression on the side lacking the filter and lower SOX2 on the filter side (Figure 7I) indicating that tissue tension impacts SOX2 expression in this progenitor domain. Moderate changes in TBXT expression were observed indicating that TBXT expression is less affected by tissue tension (Figure 7J). Thus, altering the mechanical properties of the epiblast can impact the number of SOX2/TBXT NMP cells, changing the balance of progenitors forming the growing embryonic tissues.

We next tested how tissue tension affects the lineage of mesodermal progenitors of the epiblast. We electroporated a H2B::GFP transgene in the midstreak epiblast on both sides of stage 7-8 HH embryos and placed the filter on the right side (Figure 7L). Electroporated embryos were allowed to develop overnight. Cells electroporated on the side lacking the filter ingressed to contribute to LP as expected from fate maps. In contrast, we find that electroporated cells on the filter side localize more anteriorly and integrate into the neural tube (Figure 7M-Q). Immunolabelling of these embryos shows that GFP-expressing cells can give rise to SOX2-positive cells in the neural tube on the filter side while no green cells are found at this AP location in the control side. This suggests that the electroporated cells are prevented from ingressing and remain in the epiblast eventually contributing to the neural tube. Hence changes in tissue tension can affect the lineage of cells at mid-streak level. Thus, our results demonstrate that changing the mechanical properties of the epiblast progenitors can alter their fate and identity.

## Discussion

In amniotes, the epiblast is an epithelial tissue which contains the precursors of the three embryonic germ layers. During gastrulation, cells of the epiblast ingress to form the heart, gut and notochord precursors as the PS elongates^1,2^. After it reaches its final length, the PS begins to regress, leaving in its wake the embryonic structures which organize in an anterior to posterior direction to form the future body of the embryo. At the beginning of its regression, the PS is flanked by the epiblast which contains precursors of the trunk mesodermal tissues. The distribution of these precursor territories along the AP axis reflects their future fate along the medio-lateral axis (somites, LP, blood/extraembryonic mesoderm) ^7,14,24,26^. This AP distribution also reflects the remanence of these territories in the epiblast. The extra-embryonic mesoderm precursors ingress first and thus disappear early ^22,25^, followed by LP whose precursors have fully ingressed around the 10-somite stage ^22,42^, whereas somites precursors are maintained as NMPs until the end of axis extension ^22,23^.

The multipotency of the epiblast contrasts with its well-established organization in distinct territories endowed with different fates. Our scRNAseq analysis provides evidence that during PS regression in the chicken embryo, mesodermal progenitors of the epiblast exhibit largely similar transcriptional profiles. At stage 4HH, at the beginning of PS regression, we identified a single cluster corresponding to epithelial epiblast. At stage 5HH, we started to detect a second cluster exhibiting both epiblastic and NMP characteristics. This is consistent with the first appearance of SOX2/TBXT positive cells in the anterior epiblast of the chicken embryo at this stage ^22^. These two epiblastic clusters were maintained at the 6-somite stage but at the 35-somite stage, only the NMP cluster remained. Intriguingly, no clear PS cluster was identified in our dataset, suggesting that PS and adjacent cells of the epiblast do not exhibit significant transcriptomic differences^43^. Our grafting results in the chicken embryo demonstrate that swapping epiblast cells from the LP and NMP territories results in cells adopting the fate of their new location. These results support the plasticity of epiblast cells previously reported in mouse embryos ^15^. Apart from these two epiblastic clusters, our analysis failed to identify any regionalization of epiblastic precursors that could underly established fate maps. This conclusion is consistent with recent comprehensive scRNAseq studies of gastrulating human, mouse and rabbit embryos with much higher resolution which also did not evidence epiblastic subdomains ^32,34,44–46^.

While scRNAseq analysis can be used to infer developmental trajectories, which in many cases reflect the lineage history of cells, such is not always the case. These studies therefore need to be further validated by lineage tracing. The reconstruction of cell developmental history in silico using scRNAseq data first necessitates identification of cell identities which usually relies on clustering analysis. However, defining the clustering resolution is generally highly biased by preexisting knowledge of the development of these lineages. Here, we introduce a new suite of tools to analyze developmental trajectories. A first program called Opticlust uses the significance of differentially expressed genes to define clusters in a more unbiased fashion. We also developed another module called AIntegrates, which employs machine learning to guide the annotation process. Finally, CADOT (Cluster ADjusted Optimal transport) can predict biologically relevant hypotheses regarding ancestor/descendant relationships. This new analysis pipeline aims to facilitate the prediction of cell fate transitions using scRNAseq data. Using this pipeline, we could predict ancestor/descendant hypotheses from the epiblast cell states, including both known and unknown transitions. One striking prediction of CADOT was the possibility of NMPs contributing to the LP lineage. This was unexpected as NMPs were initially reported as showing a descendance restricted to neural tube and somites ^23^. We validated this prediction using electroporation of epiblast cells to perform single-cell lineage tracing using a Brainbow-derived labeling strategy^39^. This demonstrated that single cells from the midstreak level of stage 5HH embryos can give rise to neural, mesodermal, and lateral plate descendants but they lose this capacity around stage 8HH. Surprisingly, such multipotency has not been previously reported in classical avian fate maps of the epiblast ^6,7,9–14,16,18–21,26^. However, most of the epiblast fate maps were analyzed after a 24h period, whereas our analysis was done after a much longer period of time (48-72h), unveiling previously unrecognized contributions of epiblastic territories.

To understand how the descendants of single epiblast cells can end up in distant territories such as somites, LP and neural tube, we performed a detailed analysis of cell and tissue movements at these stages. We combined in toto live imaging with the dynamic morphoskeleton pipeline ^40^, which allows to identify cell and tissue dynamics by projecting their displacements both forward and backward in time. Using PIV analysis of the epiblast from time lapse movies of chicken embryos, we could identify regions that act as attractors (towards which cells are converging) such as the PS or as repellers (from which cells are diverging) such as the boundaries between the epiblast fated to give rise to mesoderm and ectoderm. We next increased the resolution of our analysis using nuclear labeling of embryos to extract single cell tracks of epiblast cells from time lapse movies. Using these tracks with the morphoskeleton pipeline, we identified repellers corresponding to boundaries forming perpendicular to the PS and approximately delimiting known precursor domains. We first identified a B1 boundary which forms early to separate the extraembryonic and LP domains at the midstreak level of the PS. Formation of this boundary is soon followed by a B2 boundary forming at the anterior PS level to separate the LP and PM domains. This analysis suggests that prior to the formation of these boundaries, cells can move freely among these different territories of the epiblast. They become constrained to their final domain only after formation of these boundaries. We further show that prior to boundary formation, epithelial cells of the epiblast can move around in the tissue as a result of cell intercalation following cell division. This mechanism enables the displacement of daughter cells within the epiblast, allowing them to relocate into different progenitor domains. This can explain the contribution of single cells to somites, LP and NT predicted by CADOT and observed with lineage tracing ^47,48^. Such a mechanism of cell intercalation upon cell division occurs in the epiblast during earlier stages of chicken embryo gastrulation when it dissipates tissue-wide tension generated by PS formation ^49^. Cell intercalation is more frequent in the anterior epiblast than in the posterior, consistent with the higher proliferation of cells observed in this area ^22^.

We also observed a high level of anisotropic deformations dependent on cell proliferation in the anterior but not in the posterior epiblast during early stages of PS regression. This suggests that the anterior and posterior regions of the epiblast are exposed to very different mechanical constraints. To probe the mechanical environment, we monitored the deformation of alginate gels grafted in the anterior or posterior PS. We show that the anterior epiblast is exposed to significant pulling forces in the AP and ML directions while the posterior domain experiences compression associated to PS ingression. We challenged the mechanical environment of the epiblast by placing a filter on one side of the embryo in ovo. This altered the fate of epiblast cells, increasing the number of NMPs on the opposite side to the filter. Furthermore, we showed that the presence of the filter prevents normal ingression of cells of the midstreak level which contributed to the neural tube instead of LP. These experiments further demonstrate the multipotency of epiblast cells and show that mechanical constraints play a critical role in the fate determination of epiblast cells^50^.

Overall, our experiments demonstrate a striking level of multipotency of epiblast cells during early stages of PS regression. This plasticity can be attributed to relocation of the descendants of early epiblast cells to distant territories due to cell intercalation following division. Epiblast cells appear to be largely specified to their definitive fate slightly later, when movement between presumptive territories becomes restricted. The subsequent fate of these cells along the AP axis is controlled in part by differences in the mechanical environment along the epiblast.

## Acknowledgments

We thank members of the Pourquié lab, L. Mahadevan, Dan Wagner, and Cliff Tabin for their discussions and critical reading of the manuscript. We also thank Jean Livet for the MAGIC markers plasmids. We thank the NeuroTechnology Studio at Brigham and Women’s Hospital for providing the confocal LSM 880 instrument access and consultation on data acquisition and data analysis. We thank the single-cell core at Harvard Medical School and the Bauer Core facility at Harvard University for their assistance with single-cell experiments and sequencing. CG was funded by an EMBO ALTF 406-2015 fellowship, and the project was funded by NIH RO1HD097068-02 to OP. YD is the recipient of a prize from Fondation pour la Recherche Médicale (FRM) PLP2020100012456. MS was partially funded by the Hellam Fellowship.

## Authors contributions

CG conceptualized and performed most experiments under the supervision of OP. YD did the single-cell analysis and created the new algorithms and associated figures with CG. MS performed the morphoskeleton analysis from movies acquired and tracked by CG. CG, YD, and OP wrote the manuscript. OP supervised the whole project. All authors discussed the data.

## Material and methods

### Chicken and quail embryos

Fertilized specific Pathogen Free (SPF) chicken eggs were obtained from Charles River laboratories. Fertilized eggs from transgenic chickens expressing cytoplasmic GFP ubiquitously^51^ were obtained from Susan Chapman at Clemson University. Fertilized eggs from transgenic quails expressing PGK:H2B-mCherry x hUbC:Membrane-GFP were obtained from Rusty Lansford at the University of Southern California. Eggs were incubated at 38 °C in a humidified incubator, and embryos were staged according to Hamburger and Hamilton (HH)^52^ or Zacchei^53^ for chicken or quail respectively. We cultured chicken embryos mainly from stage 5HH on a ring of whatman paper on agar plates as described in the EC culture protocol^54^.

### Immunohistochemistry

For whole mount immuno-histochemistry, stage 3 to 20 HH chicken embryos were fixed in 4% paraformaldehyde (PFA)(158127, Sigma) diluted in PBS 1X at 4 °C overnight. The embryos were rinsed and permeabilized in PBS-0.1% triton, 3 times 30 min, and incubated in blocking solution (PBS-0.1% triton, 1% donkey serum (D9663, Sigma)) prior to incubating with primary and secondary antibodies. Embryos were incubated at 4 °C overnight with antibodies against TBXT/BRACHYURY (1/1000, R&D Systems: AF2085), SOX2 (1/1000, Millipore: ab5603), PHOSPHO-HISTONE 3 (1/1000, SANTA CRUZ: sc-8656), MSGN1 (1/1000)^55^, PHOSPHO-SMAD1 (1/500, Cell Signaling: D5B10) diluted in blocking solution. Embryos were rinsed and washed 3 times 30 minutes in PBS-0.1% triton, incubated 1 hour in blocking solution and incubated at 4 °C overnight with secondary antibodies conjugated with AlexaFluor (Molecular probes) diluted in blocking solution. If the staining was not imaged in the following 2 days, post fixing was performed using a 4% PFA solution.

Images were captured using a laser scanning confocal microscope with a 10X or 20X objective (LSM 780 or 980, Zeiss). To image the whole embryo, we used the tiling and stitching function of the microscope (5 by 2 matrix) and z sectioning (5mm). Later stages (from 17 HH) were imaged in clearing solution using the scale A2 clearing protocol (4 M urea 0.1% Triton, 10% glycerol)^56^. For imaging, the embryo was placed in the clearing solution 30 minutes prior to imaging in glass bottom dishes (Mattek).

### Plasmid preparation, *in ovo* electroporation for in vivo long-term tracing

#### *in ovo* electroporation

Chicken embryos at stage 5HH or 8^−^HH were prepared for in ovo electroporation. Eggs were windowed and a DNA solution (1μg/μl) mixed in HBBS, 30% glucose and 0.1% Fast-green was microinjected in the egg, in the space between the vitelline membrane and the epiblast at the 50% PS level between the B1 and B2 boundaries. Electroporation was carried out using 2 pulses at 5V for 1 msec on each side of the PS in the LP domain using a needle electrode (CUY614, Nepa Gene, Japan) and an ECM 830 electroporator (BTX Harvard Apparatus). This procedure only labels the superficial epiblast layer. Eggs were then re-incubated for further development.

#### Nucbow cell tracing

Lineage tracing was performed by co-electroporating *in ovo* the following constructs: a self-excising Cre recombinase (se-Cre), the nucbow construct and the TolII transposase as described in Loulier et al. (2014)^39^ in a 1/1/1 ratio at (1µg/µl, each). We used similar concentrations for the nucbow and transposase plasmids to that described in Loulier et al. (2014) but increased 10 times the concentration (1 µg/µl versus 0.1 µg/µl) of the se-Cre to favor fast recombination and integration. Because non-integrated nucbow plasmids can remain episomal and transiently affect the color of a cell, we performed our analyses after 36 h when the plasmids are expected to have fully diluted through cell division. 36 h after electroporation, we see that the number of fluorescent cells has significantly decreased suggesting that the episomal transgenes have now been diluted. To perform lineage analysis, we fixed the electroporated embryos at stage 17HH, and imaged them in clearing solution ScaleA2. The imaging was performed using an LSM 880 with Airyscan module in the 3 fluorescent channels using the recommended gating of Loulier et al. (2014)^39^.

#### Quantification

Cells were manually segmented in the YFP and Cherry channel using image J. Positions were assigned to the mesodermal and neural tube layers. Color retrieval was performed by measuring the intensity in the 3 channels, Cerulean, YFP and Cherry so that the total of all the intensities was normalized to 1 and expressed in percentages similarly to Loulier et al., (2014)^39^. Cluster assignment was performed using K-mean clustering followed by thresholding of cells with a silhouette >0.4. The coordinates were then calculated in a triplot diagram for visualization. Cells within the same space in the triplot have the same color coding.

### Labeling and quantification for morphometric analysis

#### Nuclear red labelling in ovo

The nuclear red solution was prepared from the NucRed™ Live 647 ReadyProbes™ Reagent (Thermofisher) and diluted in PBS. Sparse nuclear labelling of the dorsal epiblast was performed *in ovo* by injecting the nuclear red solution between the epiblast and the vitelline membrane at the PS level for 15 minutes. Embryos were then dissected, rinsed in PBS and mounted on paper filter for EC culture to perform live imaging from the dorsal side for epiblast cell tracking.

#### PIV analysis

Whole epiblast of GFP-positive embryo was analyzed by computing the velocity field using a custom version of the MATLAB PIV lab software.

#### Cell tracking analysis

Whole epiblast cell tracking was performed using the plugin from Image J. tracks were then visualized and analyzed using a custom code in Python maintained by Arthur Michaut and available here: (https://track-analyzer.pages.pasteur.fr). Cell velocities were computed by calculating the discrete displacements. In order to back-track cells in regions of interest, groups of cells were selected at any given time using selection tools provided by the Scikit-image package. Trajectories were then plotted using the Matplotlib package.

#### Dynamic Morphoskeletons

Given a planar velocity field v(x, t), we identify the Dynamic Morphoskeleton (DM)—i.e. attractors and repellers of cell motion as well as deformation maps--, from the backward and the forward Finite-Time Lyapunov Exponents (FTLE). The DM is based on a Lagrangian tissue deformation description, which combines local and global mechanisms along cell trajectories. Attractors and repellers mark embryonic regions toward which cells converge or diverge over a specific time interval t_0_, t_0_+ T (Supplementary Figure 7B). High values of the forward FTLE mark repellers; attractors are marked by high values of backward FTLE, and their domain of attraction (DOA) by high values of the backward FTLE displayed on the initial cell positions. Overall, the DM reveals the organizers of spatiotemporal trajectories and is robust to noise ^40^, hence it is ideal for quantifying morphogenesis, Repellers and deformation analysis: ^40^.

### Single cell RNA sequencing preparation and analysis

#### Preparation of single-cell suspensions for scRNA-seq

Single-cell dissociation protocols were optimized to achieve >90% viability and minimize doublets before sample collection. To generate the samples, 4 embryos were harvested for each stage and cells were dissociated and captured on an inDrops setup on the same day. Stage 4HH was added to our previous dataset including stage 5HH and 6-somite and 35 somites^22^. For new samples, we dissected 2 anterior halves of the PS including the Hensen’s Node and the posterior region of the neural plate and 2 larger regions comprising the PS and adjacent epiblast. For single cell dissociation, the dissected tissue was briefly rinsed in cold PBS, and incubated in Accutase (Gibco) for 10 min at 37 °C followed by mechanical dissociation. The cell suspension was analyzed with a hemocytometer to assess the quality of the dissociation and evaluate cell density. Dissociated cells were centrifuged at 350*g* for 5 min at 4 °C and resuspended at a concentration of 250,000 cells per microliter in 0.25% BSA in PBS. 2× 3,000 cells were sequenced per sample. Two biological replicates were collected per sample and the sequencing data from both samples were combined for data analysis.

#### Barcoding, sequencing and mapping of single-cell transcriptomes

Single-cell transcriptomes were barcoded using the inDrops pipeline using V3 sequencing adapters^30^. Following within-droplet reverse transcription, emulsions consisting of about 3000 cells were broken, frozen at −80 °C, and prepared as individual RNA-seq libraries. inDrops libraries were sequenced on an Illumina NextSeq 500 using the NextSeq 75 High Output Kits using standard Illumina sequencing primers and 61 cycles for read 1 and 14 cycles for read 2, 8 cycles each for index read 1 and index read 2. Raw sequencing data (FASTQ files) were processed using the inDrops.py bioinformatics pipeline available at https://github.com/indrops/indrops. Transcriptome libraries were mapped to *Gallus gallus* transcriptome built from the GRCg6a (GCA_000002315.5) genome assembly. Bowtie version 1.1.1 was used with parameter –e 200.

#### Processing of scRNA-seq data

Single-cell counts matrices were processed and analyzed using ScanPy (1.4.3) and custom Python scripts (Code Availability). Low-complexity cell barcodes, which can arise from droplets that lack a cell but contain background RNA, were filtered in two ways. First, inDrops data were initially filtered to only include transcript counts originating from abundantly sampled cell barcodes. This determination was performed by inspecting a weighted histogram of unique molecular identifier–gene pair counts for each cell barcode, and manually thresholding to include the largest mode of the distribution. Second, low-complexity transcriptomes were filtered out by excluding cell barcodes associated with <400 expressed genes. Transcript unique molecular identifier counts for each biological sample were then reported as a transcript × cell table, adjusted by a total-count normalization, log-normalized, and scaled to unit variance and zero mean. Unless otherwise noted, each dataset was subset to the 1,000 most highly variable genes, as determined by a bin-normalized overdispersion metric.

#### Low-dimensional embedding and clustering

Processed single-cell data were projected into a 50-dimensional PCA subspace, (*k* = 10 except 35 somite k=15) nearest-neighbor graph using Euclidean distance and 50 PCA dimensions and visualized using UMAP (Uniform Manifold Approximation and Projection) representation. Clustering was performed using Louvain or Leiden community detection algorithms.

#### Identification of differentially expressed genes

Transcripts with significant cluster-specific enrichment were identified by t-test comparing cells of each cluster to cells from all other clusters in the same dataset. Genes were considered differentially expressed if they met the following criteria: log-transformed fold change > 0, adjusted *P* value < 0.05. False discovery rate (FDR) correction for multiple hypothesis testing was performed as described using Benjamini–Hochberg. The differentially expressed genes, ranked by FDR-adjusted *P* values, associated fold changes, and sample sizes (number of cells per cluster) are reported in Extended data Table1.

#### Opticlust

We develop an unbiased clustering method allowing the optimal clustering of cell types from a scRNAseq dataset. This method, called OptiClust considers the significance of the genetic profile of each identified cell population available in the scRNA sequencing data. This program was created to find the optimal cluster resolution possible according to the significance of all the genes for each cluster. OptiClust starts by defining clusters at the lowest resolution possible and increase the resolution number by a pre-defined step until 2 clusters are defined using Leiden. Once 2 clusters are defined, for a given clustering resolution, the program checks the highest adjusted p-value of the top 1 DEG sorted by score for each cluster. OptiClust consciously increase the resolution at every passage by the pre-defined step until it reaches the highest clustering resolution value where the adjusted p-value of one of the first top genes of one of the clusters is inferior to 0.05. Adjusted p-value was obtained using wilcoxon with Benjamini-Hochberg correction method. This resolution called “optimal value” is then stored in the adata and returned to the user. A visualization tool is also provided in the form of a python widget to automatically display the given resolution and the DEG for every clusters (all the computed leiden resolutions and DEG are also stored and instantly available for the user). OptiClust also include other parameters for fine tuning: position of the gene that needs to be significant in the ranked gene list, resolution range and incremental steps that needs to be tested, p-value adjusted to test, tie-correction and correction-method.

#### CADOT

We developed a tool called CADOT (Cluster ADJusted Optimal Transport) that use probability vectors for every cell to be on the trajectory of a specific clusters at the last timepoint of a multi-timepoint dataset (with at least 3 timepoints) and transition matrices inside a dataset to display the transitions from clusters to clusters between every timepoints. The probability values and the transitions matrices are obtained using Waddington-OT (WOT) 1.0.7 conceptual framework)^38^. WOT allows us to infer the temporal couplings of cells from the different samples collected independently at various timepoints and get transport matrices. Every timepoints are subsampled to contain to contain the same number of randomly selected cells. Transport matrices (also called ‘transport maps’) are created by connecting each pair of timepoints and using an estimate of cellular growth rates (to estimate the growth rate, each cell are scored according to its expression of various gene signatures like proliferation and apoptosis as described in Schiebinger et al. (2019)^38^ and then model cellular growth with a Birth-Death Process, which assigns each cell a rate of division and a rate of death). Trajectory refers to the sequence of ancestor distributions at earlier time points and descendant distributions at later time points of a cell set C. In the case of CADOT, ancestors’ probability was used and calculated by pushing back through the transport map. Transition matrices are calculated to show the amount of mass transported from a cell type to another from a start and an end point. The following WOT parameters are used: ε = 0.05 (controls the degree of entropy in the transport map), λ1 = 1 and λ2 = 50 (A smaller value of λ1 or larger value of λ2 enforces the constraints less strictly, which is useful when we do not have precise information about the growth rate). CADOT requires to use a clustering technique to define cell populations and calculate means and standard deviations of probabilities among cells from the same clusters (Given a set C of cells at time j, we use the probability to be an ancestor of C at an earlier time point i < j). Then, confidence intervals can be calculated for every cluster and the lowest confidence interval value for all cluster transitions are attributed to every cluster to correct the mean probability (this allows us to correct clusters with small numbers of outlier cells). Filtering clusters probabilities and the transition matrix by user-defined quantiles allow us to visualize the different transitions according to their levels of probability and filter background noise. The result was summarized in a directed data frame containing clusters as node, transitions as edges, probability of transition as edge colors and mass transported as edge sizes. A scanpy compatible python package, containing a WOT wrapper and CADOT computation and visualization functions is available.

### Epiblast Grafting

Donor GFP-positive embryos were isolated using the filter method in HBSS and placed on a dissecting plate with a black background. The donor embryo was turned ventral side up, endodermal and mesodermal layers were peeled away and a piece of epiblast (∼20 to 50 cells) was cut out using home-made scalpels. Each epiblast piece was checked to ensure that no mesodermal/endodermal cells remained attached before grafting. A slit in the non-GFP host at the desired location was made to place the donor epiblast. For Lateral plate domain grafts, the epiblast was taken at the 50% mark of the Primitive streak, while the NMP grafts were taken from the epiblast right below the node. We usually took an NMP and an LP epiblast graft from the same GFP donor embryo. Upon regrafting, we removed the excess HBSS and checked that the graft was well inserted into the desired region before putting the embryos back into the incubator for further development on plate culture media. In the majority of cases, the initial graft was cut asymmetrically to ensure proper AP positioning upon regrafting. No differences between asymmetrical and non-asymmetrical grafts were observed indicating that AP direction did not affect the graft results within such tiny epiblast pieces.

### Surgeries for Gel Implants and Quantification

Stage HH5 GFP-positive embryos were used for surgery experiments. Surgeries were performed *in ovo* under a Leica M90 dissecting scope using a NIGHTSEA GFP lamp in a homemade incubation chamber. Cutting was performed with a homemade scalpel from the dorsal side. The vitelline membrane over the surgery site was first slit in the middle and gently peeled on either side to make a triangular shape opening. Square regions were removed in the dorsal epiblast in the 90% or 50% PS regions. For gel implant, the cleared cut opening received an injection of 1% (w/v) alginic acid sodium (Sigma Aldrich) solution. Injected alginic solution formed gel with calcium ion in the embryo culture and integrated much better than transplant of pre-formed gel. The eggs were then re-incubated, and pictures were taken every hour using a zeiss axiocam MRC camera. Here, note that the cut was made only in the epiblast to make sure that the gel stays on top of the endodermal layer. In case the endoderm was also cut, the gel sinked into the yolk and the embryo was discarded. To quantify the evolution of gel implants, we used the Image J software to draw the contour of the graft at different times and retrieve the area. We also measure the AP and ML length of the gel. We then plotted the evolution of the area and the axes over time using Excel software by normalizing to the initial size.

### Surgery for filter experiment to hamper tissue flow

Stage HH8 embryos were used for filter surgery experiments. Surgeries were performed *in ovo* under a Leica M90 dissecting scope in a homemade incubation chamber. A first ring of Whatman paper was placed on top of the vitelline membrane with the embryo in its center and not covered by the filter. The vitelline membrane over the surgery site was first slit in the middle and gently peeled to uncover the embryo. A second rectangular-shaped Whatman filter was placed parallel to the AP axis along the PS on the right side. Eggs were then re-incubated for 6 or 12h and pictures were taken at the beginning of the experiment using a zeiss axiocam MRC camera. Following filter surgery, part of the embryos were electroporated with H2B::GFP plasmid at the 50% PS level using 1 pulse on each side of the PS and a 100 μm needle electrode. At the end of incubation, embryos were retrieved, fixed, and immunostained using the standard methods described above.

To measure tissue relaxation after cutting along the PS with and without the filter, we used GFP-positive embryos, an performed video recording under a Leica M90 dissecting scope with a zeiss axiocam MRC camera piloted by micromanager software and a NIGHTSEA GFP lamp. Relaxation was measured using the kymograph module of FIJI software image analysis.

## Supplementary Figure titles and legends

**Supplementary Figure 1:**
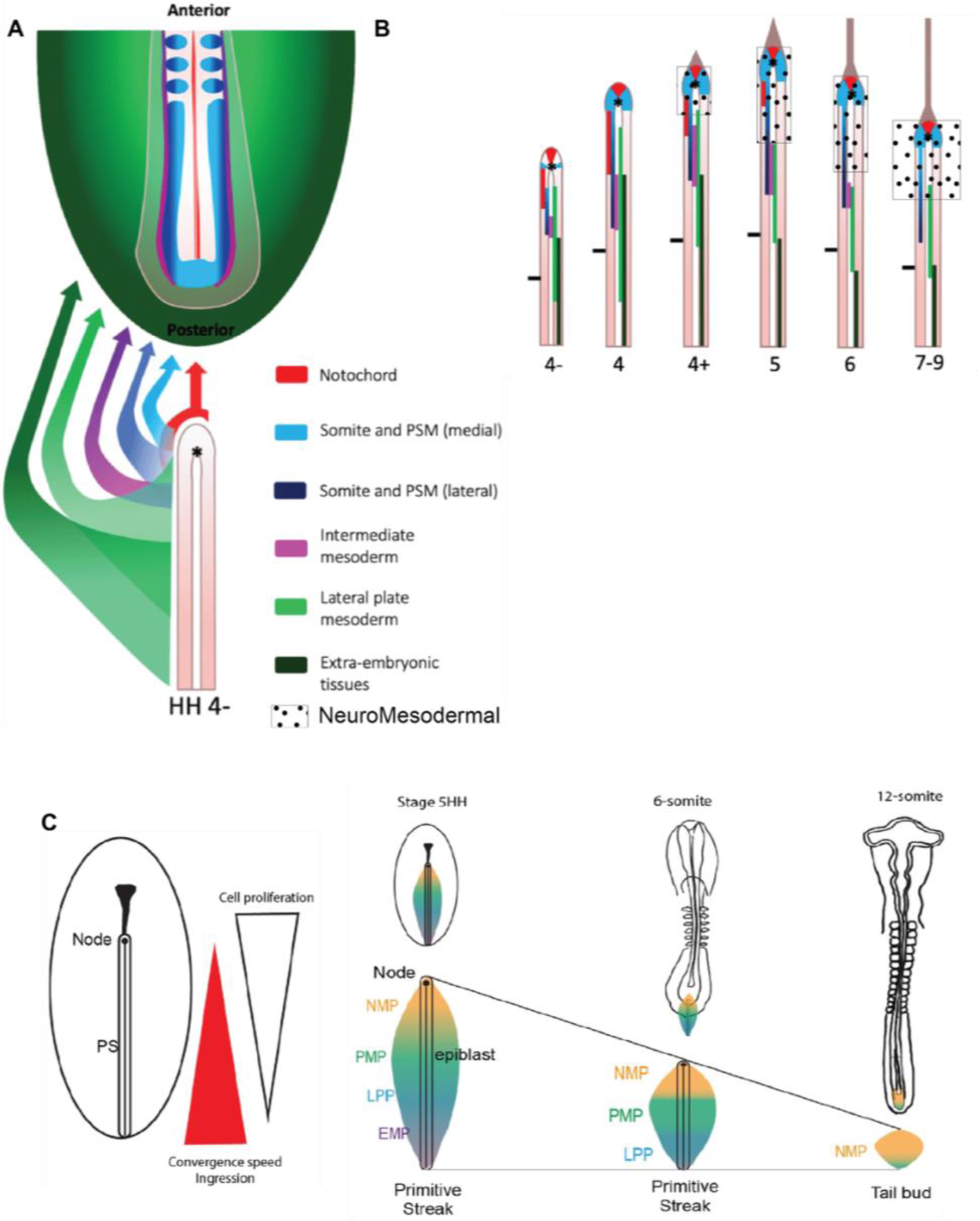
Organization of the fate, dynamics, and signaling of trunk epiblastic mesodermal precursors of chicken embryos. (A) Fate map showing the contribution of the epiblast from different AP levels to mesodermal tissues during PS regression in the chicken embryo (adapted from ^7^). (B) Position of the presumptive territories of the epiblast and the localization of the NMP domain (dots) during early stages of PS regression (between stage 4-HH to 7-9HH) (adapted from ^7^). (C) Diagram showing the gradient of cell movements along the PS during mesoderm ingression (adapted from ^22^). NMP: Neuromesodermal, PMP: Paraxial Mesoderm Progenitors; LPP: Lateral Plate Progenitors; EMP: Extra Embryonic Progenitors.

**Supplementary Figure 2:**
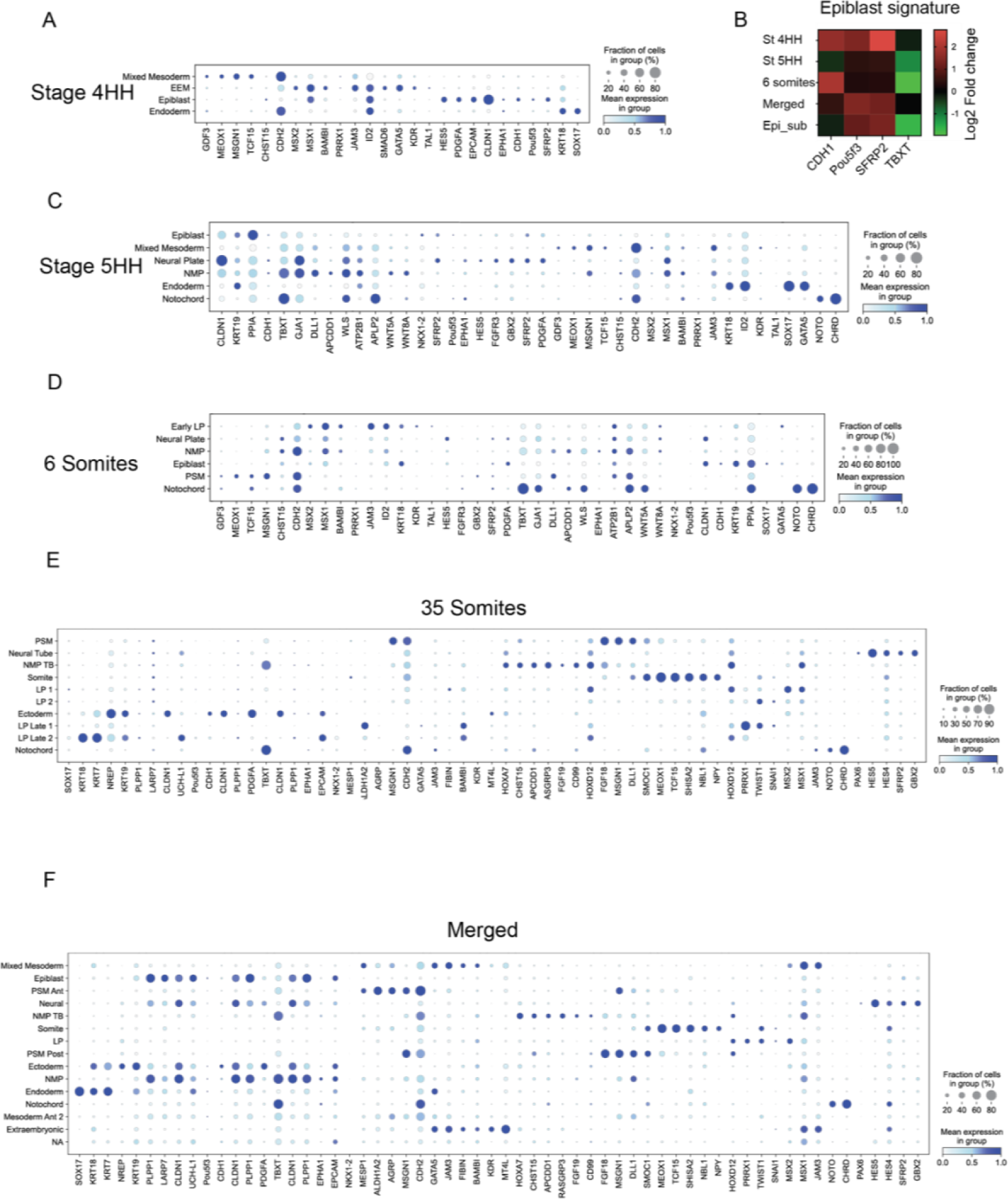
Identification of cell clusters based on marker genes’ expression. (A) Dot plots showing the expression of the marker genes used to identify the cluster identity at stage 4HH (B) Heat map showing the expression in log2 fold change of epiblast markers genes in the different epiblast clusters of 4HH, 5HH, 6 somites from Figure 2D merged and subclustered. Epi_sub corresponds to the epiblast cluster shown in Supplementary Figure 4C. (C to F) Dot plots showing the expression of the marker genes used to identify clusters at stage 5 HH (C), 6 somites (D), 35 somites (E), and all the stages merged together (F).

**Supplementary Figure 3:**
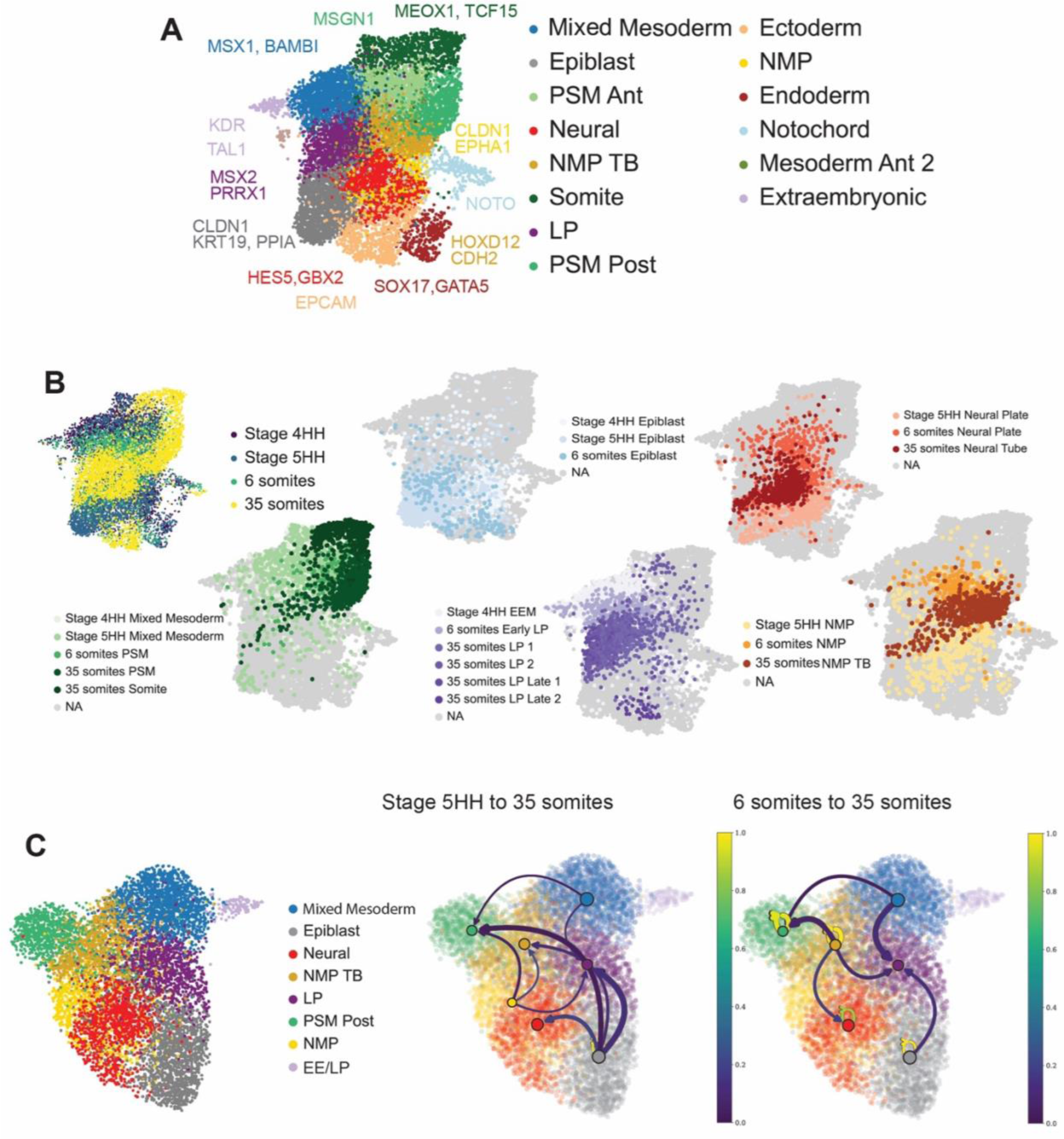
CADOT and AIntegrate: trajectory and annotations methods. (A). UMAP embedding of HH4, HH5, 6 somites, 35 somites chicken embryos merged and processed using bkknn batch correction (50 PC dimensions, 12,000 cells). Colors indicate cell-type annotations of each independent clusters specific to each timepoints, accuracy of annotations was checked using AIntegrate. (B). UMAP embedding of HH4, HH5, 6 somites, 35 somites chicken embryos merged and processed using bkknn batch correction (50 PC dimensions, 12,000 cells). Colors show the timepoints (upper left image) and the projected space of the cell-type annotations (all other images). (C). (Left) UMAP embedding of the clusters of interest (in C) for trajectory analysis. Colors indicate cell-type annotations of each independent clusters specific to each timepoints. (Right). CADOT analysis of cell movement (cluster to cluster transitions) from stage 5HH and 6 somites to stage 35 somites. Arrow colors indicate the probability of the transition, arrow size indicates the associated quantity of cells transported from one cluster to another in the merged dataset containing stage 4HH, 5HH, 6 somites, and 35 somites on clusters of interests.

**Supplementary Figure 4:**
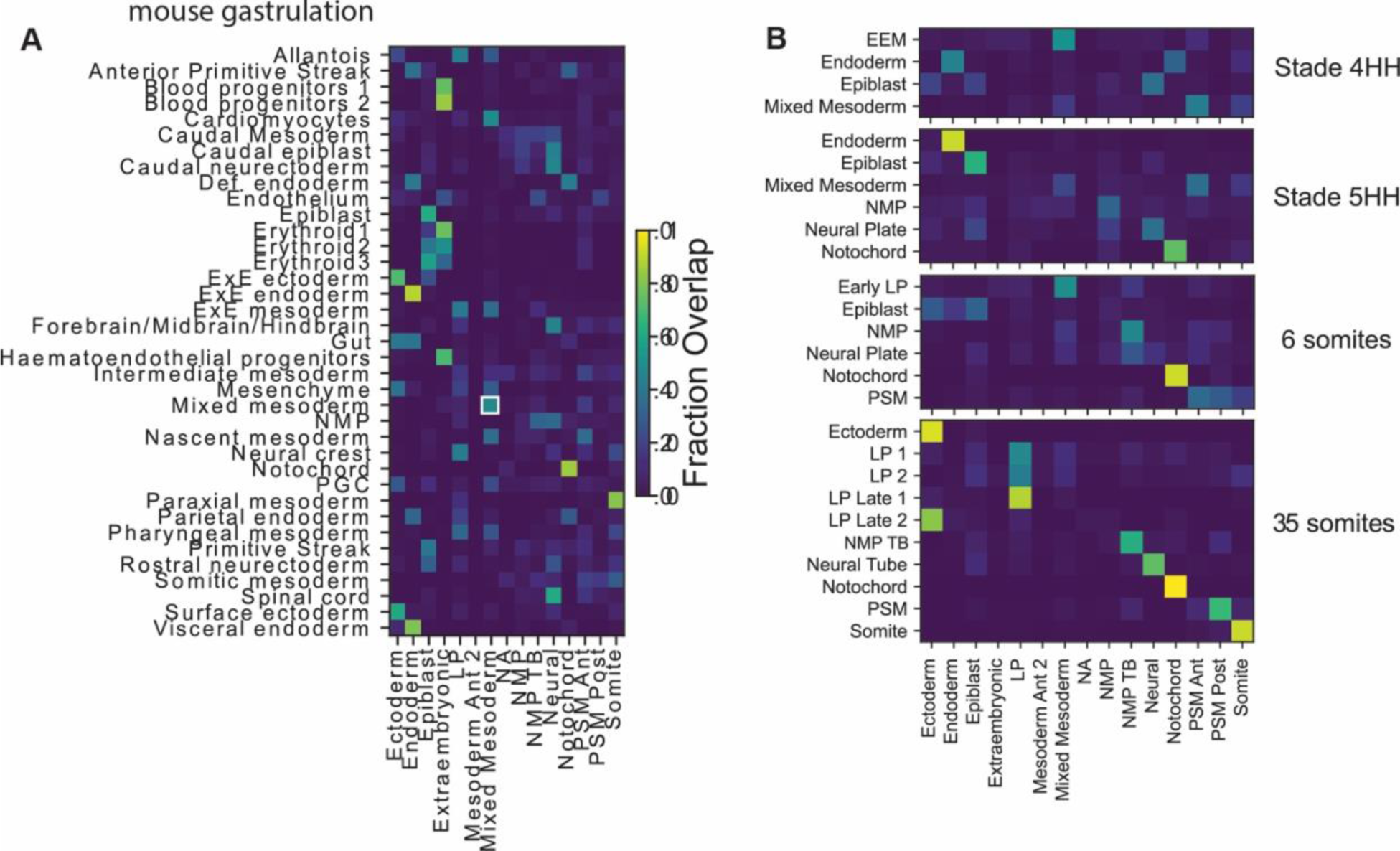
Classifier to assign cluster identity using AIntegrate. (A) AIntegrate classifier showing the correlation (fraction overlap) between the merged chicken dataset from this study and the mouse gastrulation clustering from the Pijuan-Sala (2017) study (left). (B) AIntegrate classifier showing the correlation (fraction overlap) between annotations in the integrated and merged dataset containing stage 4HH, 5HH, 6 somites and 35 somites and individual datasets annotated using the Opticlust clustering method (right).

**Supplementary Figure 5:**
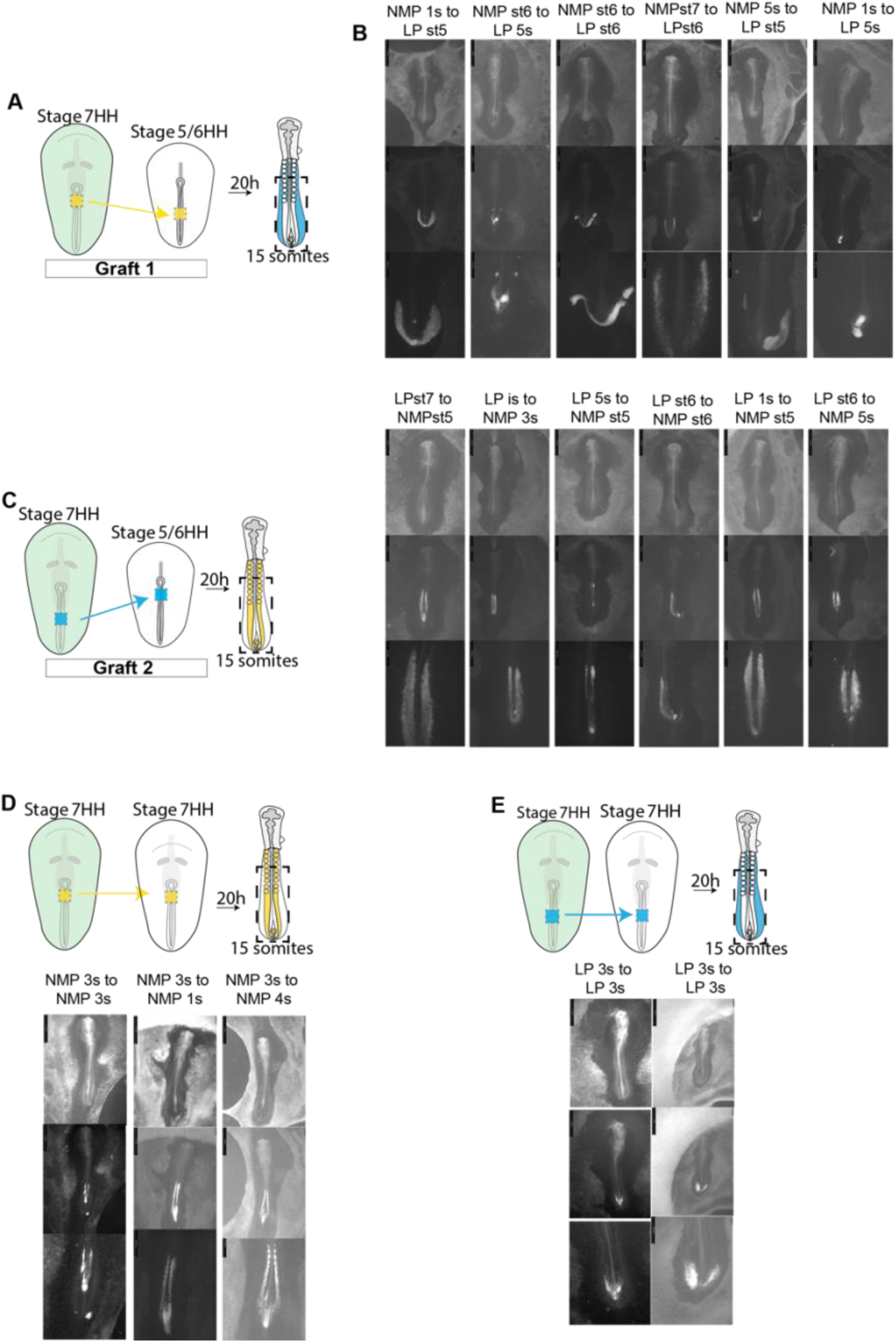
Testing developmental potencies of the epiblast. (A to C) Schemes of the graft experiments testing the developmental potency of cells of the predictive NMP (A, type 1 grafts) and LP (B, type 2 grafts) progenitors’ domains of the epiblast using transgenic donor embryos expressing ubiquitous GFP and wild type (GFP negative) hosts. (B) (Top), bright field images of grafted embryos. (Middle) lower, (bottom) higher magnifications showing the donor derived GFP expressing cells in the grafted embryos. (D) (Top) Diagram showing the homotopic and isochronic graft of a stage 7-8 HH fragment of epiblast of the NMP progenitor domain into the NMP progenitor domain of stage 7-8 HH host (control). (Top), bright field images of grafted embryos. (Middle) lower, (bottom) higher magnifications showing the donor derived GFP expressing cells in the grafted embryos. (E) (Top) Diagram showing the homotopic and isochronic graft of a fragment of the epiblast of the LP epiblast progenitor domain of stage 7-8 HH donor into the LP progenitor domain of stage 7-8 HH host (control grafts). (Top), bright field images of grafted embryos after overnight incubation. (Middle) lower, (bottom) higher magnifications showing the donor derived GFP expressing cells in the grafted embryos.

**Supplementary Figure 6:**
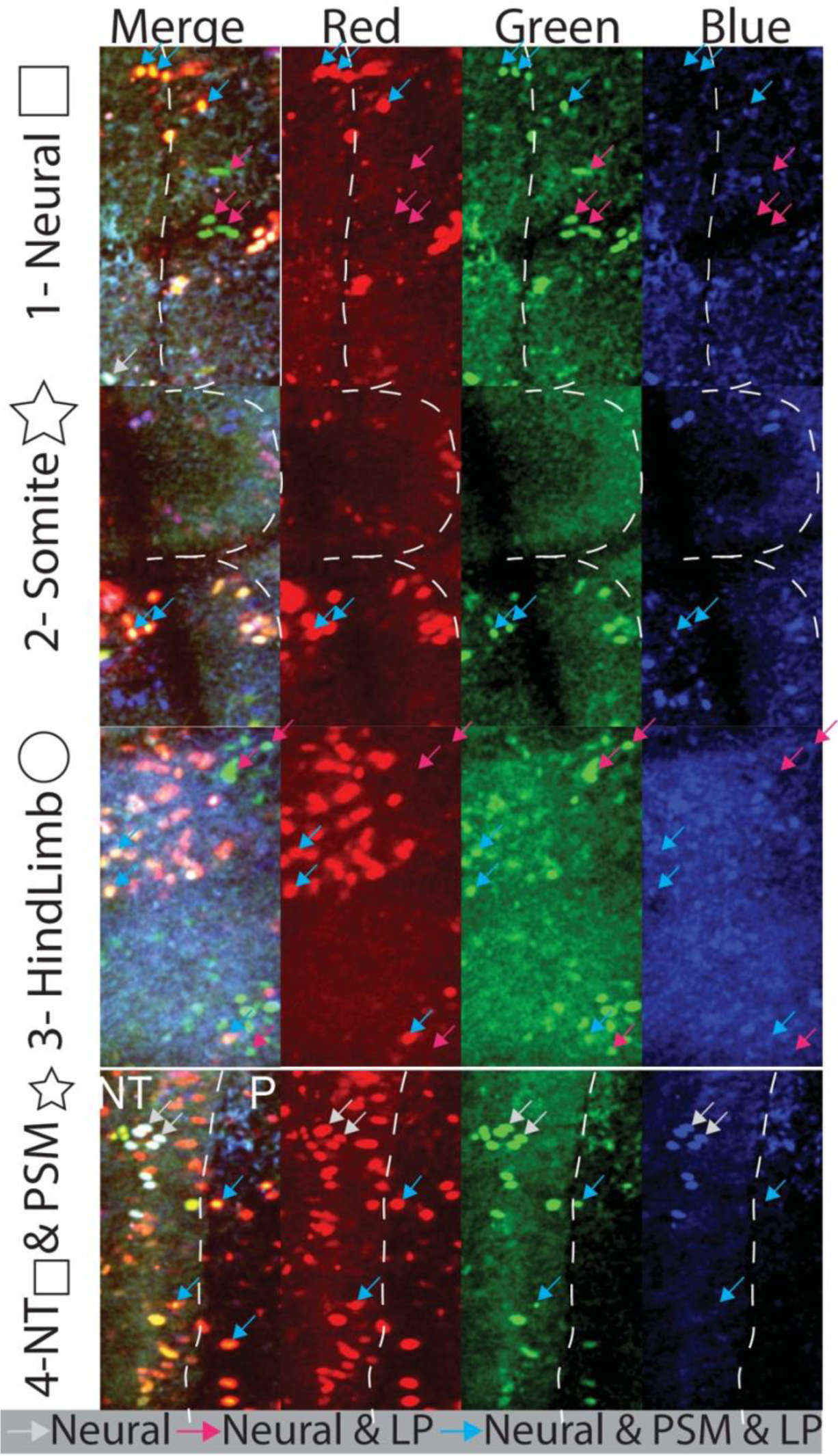
Single cell lineage tracing of the epiblast of the 50% primitive streak level at stage 5HH. Higher magnification of the regions in orange boxes in Figure 3B showing color-coded cells with different fates. Grey arrows show a representative clone of only neural cells with the same color code: 30/30/30 Red/Green/Blue corresponding to the clone 7 yellow of Figure 3D; red arrows show a representative clone of Neural and Lateral plate mesodermal cells with a color code 10/80/10 Red/Green/Blue, corresponding to the clone 11 purple of Figure 3D and light blue arrows show cells in the Neural, Paraxial and Lateral plate cells with 65/25/10 Red/green/Blue color code corresponding to the clone 3 light blue of Figure 3 D. Square: neural cells, star: PSM cell, Circle: Lateral plate cell.

**Supplementary Figure 7:**
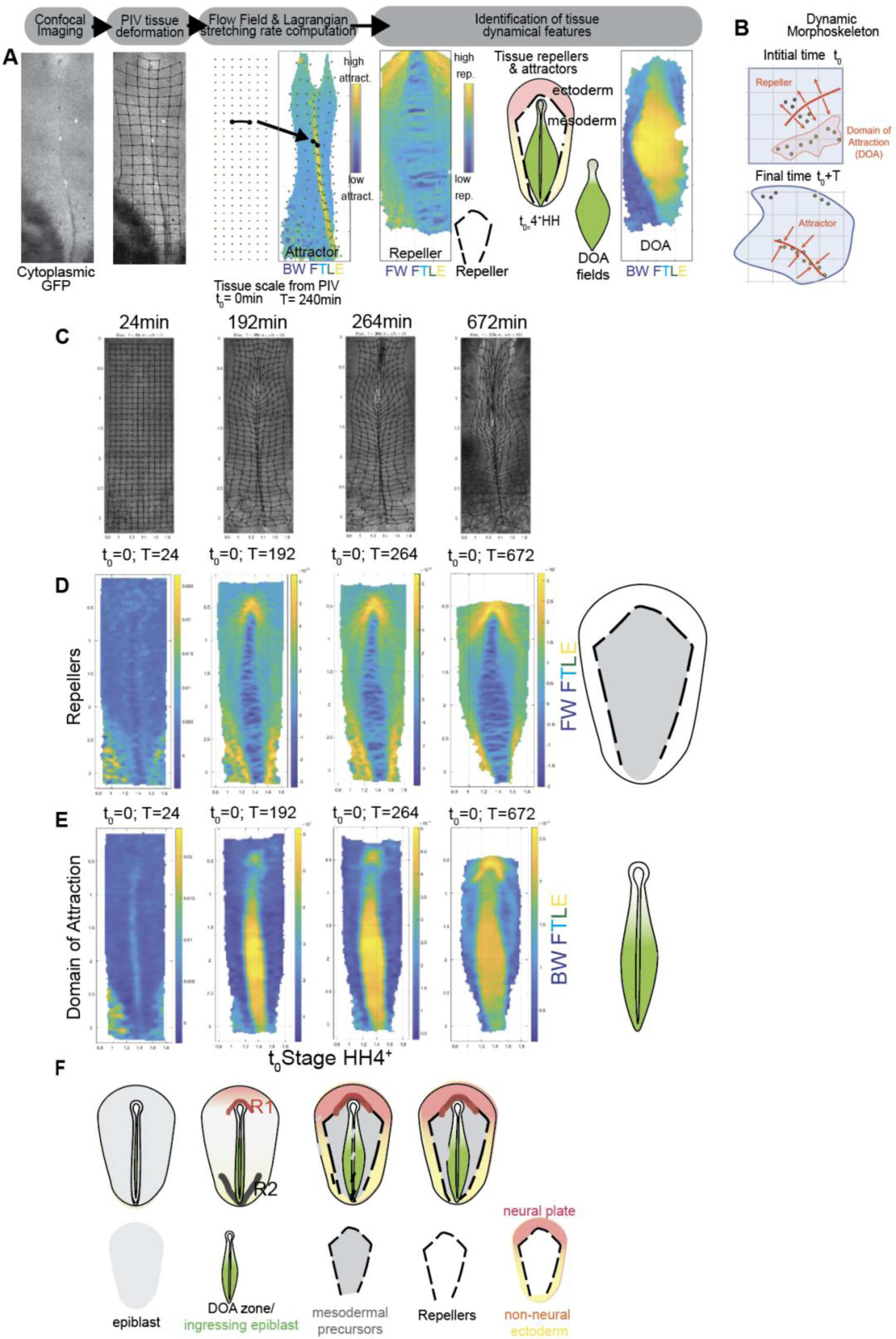
Dynamic morphoskeleton analysis using particle image velocimetry (PIV) during early stages of PS regression. (A) Workflow to identify repellers and attractors using PIV analysis and the morphoskeleton pipeline during early stages of PS regression. Representative confocal image of cytoplasmic GFP embryos used to calculate grid deformation and flow field used to compute the Forward (FW) and Backward (BW) FTLE to respectively identify the repellers and attractors as summarized in the diagram. (B) A repeller marks a curve at the initial (t_0_) embryo configuration, whose cells on its opposite sides will separate maximally by the final time (t_0_ + T). An attractor marks a curve at the final embryo configuration towards which initially distant cells will converge by (t_0_ + T). The Domain of Attraction (DOA) marks the initial embryonic region that will converge to the attractor. (C) Images of GFP-expressing transgenic embryos spanning 12 hours from the beginning of PS regression. The over-imposed grid visualizes tissue deformations (grid size 80μm). (D and E) Representative images (left) and diagrams (right) of the cytoplasmic GFP embryos analyzed with the dynamic morphoskeleton pipeline showing the repellers (D) and DOA (E) regions in WT embryos (n=3). (F) Schematics representing the position of the identified repellers (R) and DOA relative to the different precursors’ territories in a stage 4HH embryo.

**Supplementary Figure 8:**
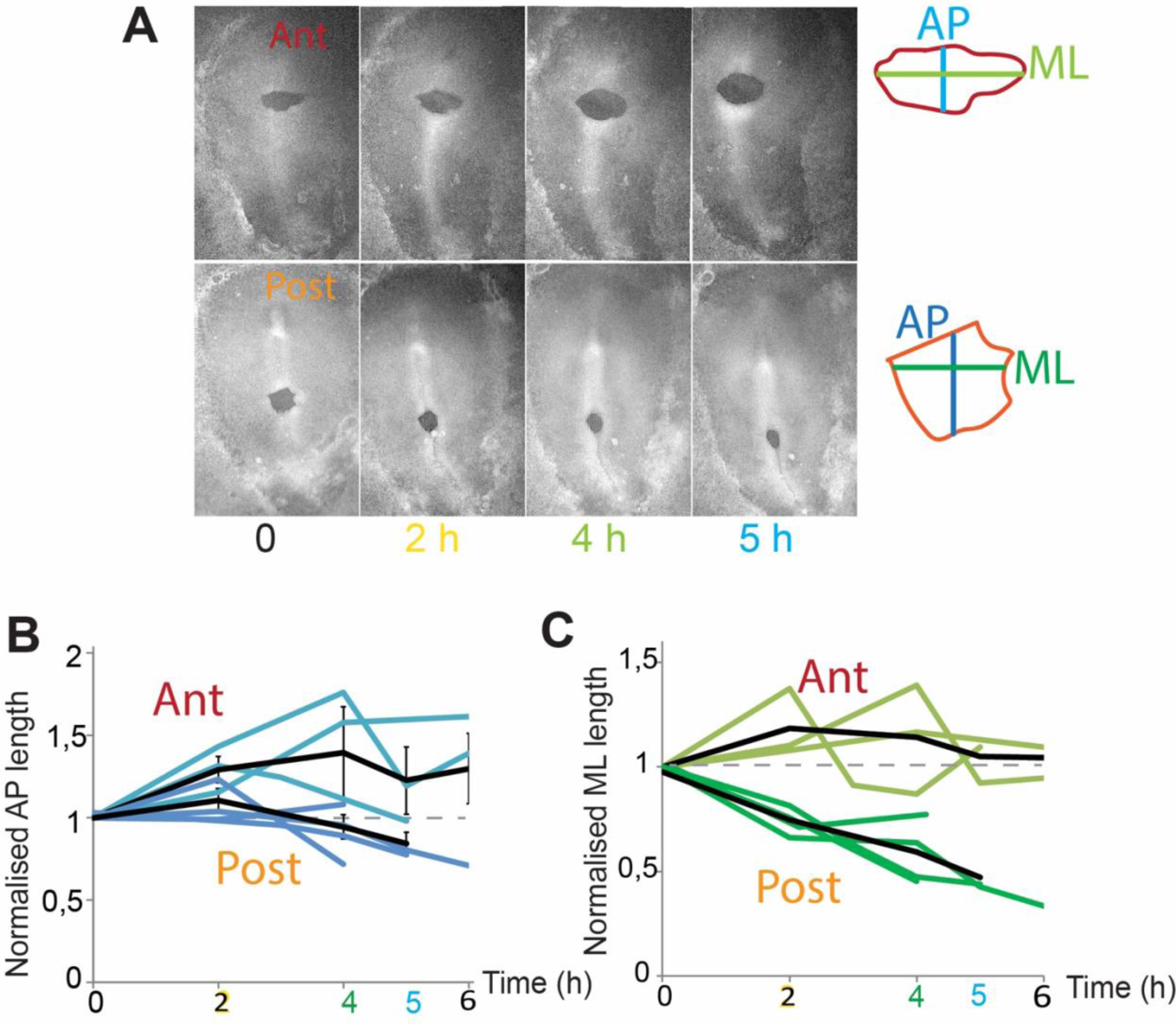
Analysis of the mechanical properties of the anterior and posterior PS. Images (A) and quantifications (B, C) of the AP (B) and ML (C) normalized length of alginate gels implanted in the anterior and posterior PS regions in ovo and analyzed every hour for 5 h. (n=8).

**Supplementary Figure 9:**
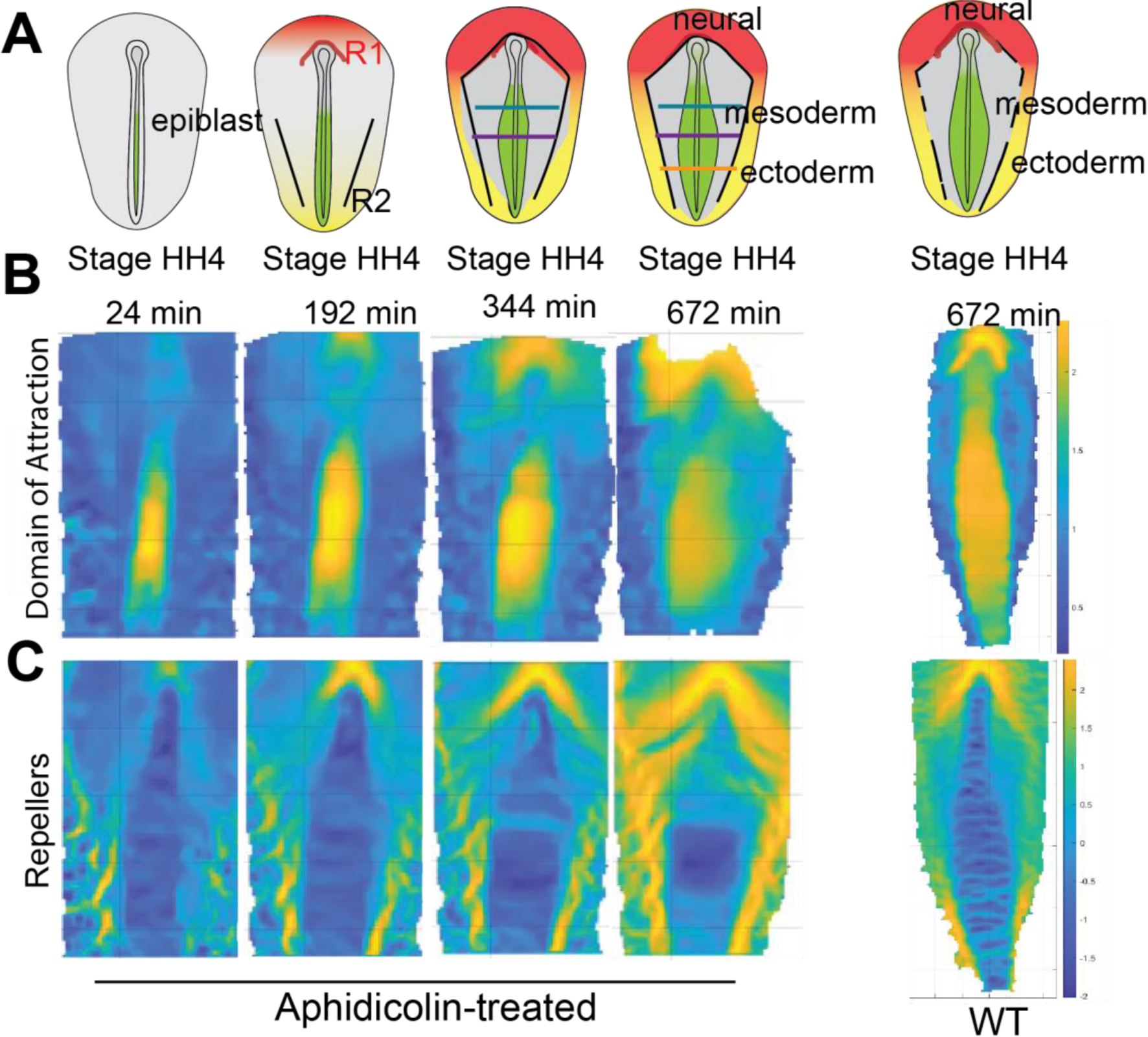
Identification of the repellers and attractors during PS regression in aphidicolin-treated embryos. (A) Diagrams and representative images of cytoplasmic GFP embryos analyzed by PIV and the morphoskeleton pipeline showing the DOA (B) and repellers (C) regions in the epiblast of aphidicolin-treated embryos (left) compared to WT embryos (right) during early stages of PS regression (n=3).

**Supplementary Figure 10:**
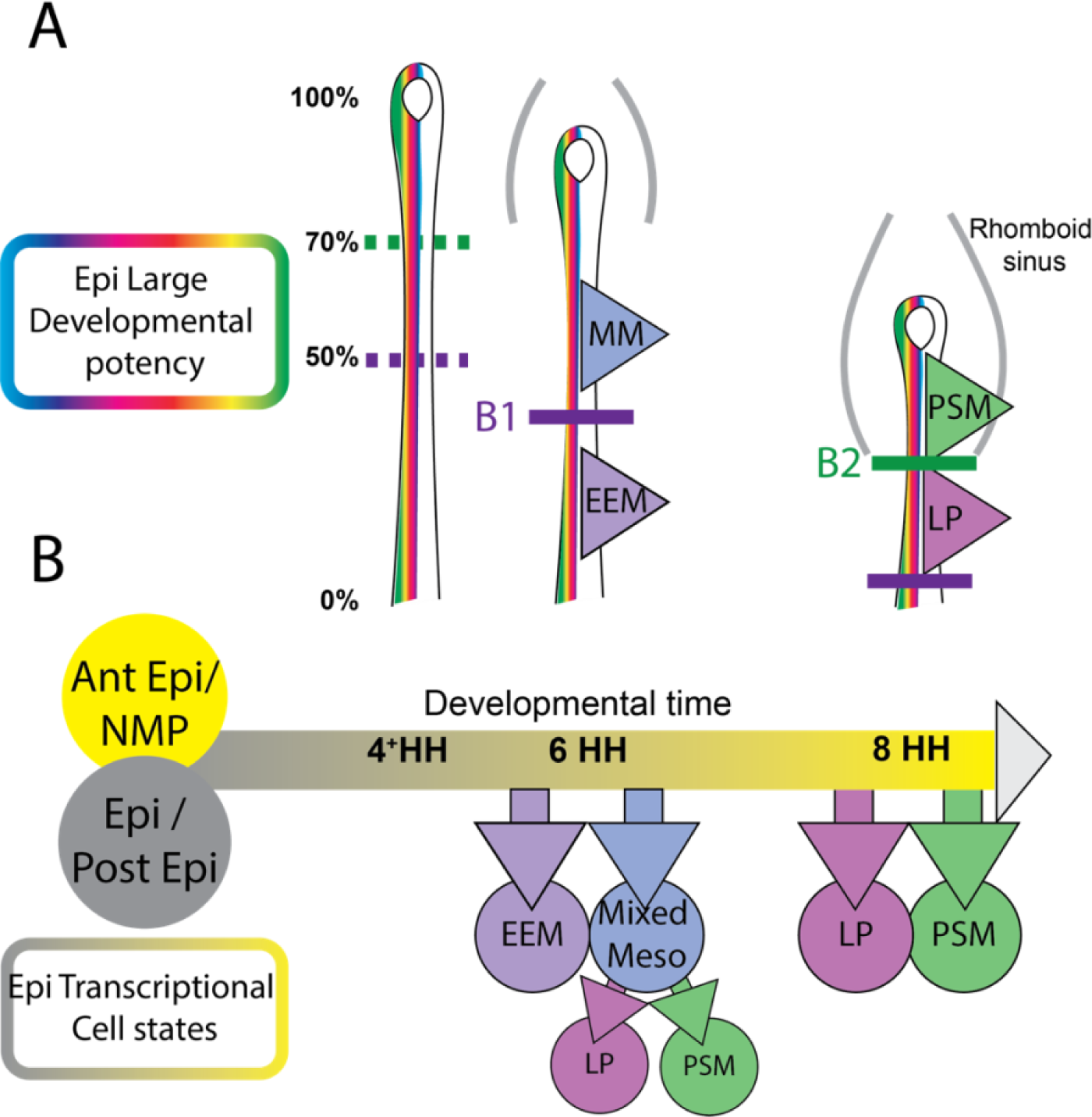
Dynamic fate map of epiblast mesoderm progenitors during PS regression in chicken embryos. (A) Diagram showing the progressive formation of boundaries between presumptive mesoderm territories of the epiblast during PS regression. (B) Diagram showing the temporal sequence of mesodermal fate diversification from the two epiblast cell states (yellow and grey) with large developmental potency (rainbow). MM: Mixed mesoderm, EEM: Extraembryonic mesoderm, LP: Lateral Plate mesoderm, PSM: Paraxial mesoderm. B1, B2: Boundary 1 and 2.

## References

1. Stern, C.D. (2004). Gastrulation in the chick. In Gastrulation: from cells to embryos, C.D. Stern, ed. (Cold Spring Harbor Laboratory Press), pp. 219–232.

2. Sheng, G., Martinez Arias, A., and Sutherland, A. (2021). The primitive streak and cellular principles of building an amniote body through gastrulation. Science 374, abg1727. 10.1126/science.abg1727.

3. Pasteels, J. (1937). Etudes sur la gastrulation des Vertébrés méroblastiques. III. Oiseaux. IV. Conclusions générales. Arch.Biol. 48, 381–488.

4. Gont, L.K., Steinbeisser, H., Blumberg, B., and de Robertis, E.M. (1993). Tail formation as a continuation of gastrulation: the multiple cell populations of the Xenopus tailbud derive from the late blastopore lip. Development 119, 991–1004.

5. Catala, M., Teillet, M.A., and Le Douarin, N.M. (1995). Organization and development of the tail bud analyzed with the quail-chick chimaera system. Mech.Dev 51, 51–65.

6. Hatada, Y., and Stern, C.D. (1994). A fate map of the epiblast of the early chick embryo. Development. 120, 2879–2889.

7. Psychoyos, D., and Stern, C.D. (1996). Fates and migratory routes of primitive streak cells in the chick embryo. Development. 122, 1523–1534.

8. Tam, P.P., and Behringer, R.R. (1997). Mouse gastrulation: the formation of a mammalian body plan. Mech Dev 68, 3–25.

9. Charrier, J.B., Teillet, M.A., Lapointe, F., and Le Douarin, N.M. (1999). Defining subregions of Hensen’s node essential for caudalward movement, midline development and cell survival. Development 126, 4771–4783.

10. Catala, M., Teillet, M.A., De Robertis, E.M., and Le Douarin, M.L. (1996). A spinal cord fate map in the avian embryo: while regressing, Hensen’s node lays down the notochord and floor plate thus joining the spinal cord lateral walls. Development. 122, 2599–2610.

11. Rosenquist, G.C. (1966). A radioautographic study of labeled grafts in the chick blastoderm. Development from primitive-streak stages to stage 12. Contr. Embryol. Carnegie Inst. Wash. 38, 71–110.

12. Nicolet, G. (1970). Analyse autoradiographique de la localisation des differentes ebauches presomptives dans la ligne primitive de l’embryon de poulet. J. Embryol. Exp. Morph. 23, 79–108.

13. Nicolet, G. (1971). Avian gastrulation. Adv Morphog 9, 231–262.

14. Iimura, T., Yang, X., Weijer, C.J., and Pourquie, O. (2007). Dual mode of paraxial mesoderm formation during chick gastrulation. Proc Natl Acad Sci USA 104, 2744–2749.

15. 15. Wymeersch, F.J., Huang, Y., Blin, G., Cambray, N., Wilkie, R., Wong, F.C., and Wilson, V. (2016). Position-dependent plasticity of distinct progenitor types in the primitive streak. eLife 5. 10.7554/eLife.10042.

16. Cambray, N., and Wilson, V. (2002). Axial progenitors with extensive potency are localised to the mouse chordoneural hinge. Development 129, 4855–4866.

17. Cambray, N., and Wilson, V. (2007). Two distinct sources for a population of maturing axial progenitors. Development 134, 2829–2840.

18. Garcia-Martinez, V., Alvarez, I.S., and Schoenwolf, G.C. (1993). Locations of the ectodermal and nonectodermal subdivisions of the epiblast at stages 3 and 4 of avian gastrulation and neurulation. J Exp Zool 267, 431–446.

19. Brown, J.M., and Storey, K.G. (2000). A region of the vertebrate neural plate in which neighbouring cells can adopt neural or epidermal fates. Curr Biol 10, 869–872.

20. Selleck, M.A., and Stern, C.D. (1991). Fate mapping and cell lineage analysis of Hensen’s node in the chick embryo. Development. 112, 615–626.

21. Spratt Jr., N.T. (1955). Analysis of the organizer center in the early chick embryo. I. Localisation of prospective notochord and somite cells. J.Exp.Zool. 128, 121–162.

22. Guillot, C., Djeffal, Y., Michaut, A., Rabe, B., and Pourquie, O. (2021). Dynamics of primitive streak regression controls the fate of neuromesodermal progenitors in the chicken embryo. eLife 10. 10.7554/eLife.64819.

23. Tzouanacou, E., Wegener, A., Wymeersch, F.J., Wilson, V., and Nicolas, J.F. (2009). Redefining the progression of lineage segregations during mammalian embryogenesis by clonal analysis. Dev Cell 17, 365–376. 10.1016/j.devcel.2009.08.002.

24. Tam, P.P., and Beddington, R.S. (1987). The formation of mesodermal tissues in the mouse embryo during gastrulation and early organogenesis. Development 99, 109–126.

25. Spratt Jr., N.T. (1947). Regression and shortening of the primitive streak in the explanted chick blastoderm. J Exp Zool 104, 69–100.

26. Schoenwolf, G.C., Garcia-Martinez, V., and Dias, M.S. (1992). Mesoderm movement and fate during avian gastrulation and neurulation. Dev Dyn. 193, 235–248.

27. Selleck, M.A., and Stern, C.D. (1992). Commitment of mesoderm cells in Hensen’s node of the chick embryo to notocord and somite. Development 114, 403–415.

28. Tonegawa, A., and Takahashi, Y. (1998). Somitogenesis controlled by Noggin. Dev Biol. 202, 172–182.

29. Streit, A., and Stern, C.D. (1999). Mesoderm patterning and somite formation during node regression: differential effects of chordin and noggin. Mech Dev 85, 85–96.

30. Klein, A.M., Mazutis, L., Akartuna, I., Tallapragada, N., Veres, A., Li, V., Peshkin, L., Weitz, D.A., and Kirschner, M.W. (2015). Droplet barcoding for single-cell transcriptomics applied to embryonic stem cells. Cell 161, 1187–1201. 10.1016/j.cell.2015.04.044.

31. Onichtchouk, D. (2016). Evolution and functions of Oct4 homologs in non-mammalian vertebrates. Biochim Biophys Acta 1859, 770–779. 10.1016/j.bbagrm.2016.03.013.

32. Tyser, R.C.V., Mahammadov, E., Nakanoh, S., Vallier, L., Scialdone, A., and Srinivas, S. (2021). Single-cell transcriptomic characterization of a gastrulating human embryo. Nature 600, 285–289. 10.1038/s41586-021-04158-y.

33. Guibentif, C., Griffiths, J.A., Imaz-Rosshandler, I., Ghazanfar, S., Nichols, J., Wilson, V., Gottgens, B., and Marioni, J.C. (2021). Diverse Routes toward Early Somites in the Mouse Embryo. Dev Cell 56, 141–153 e146. 10.1016/j.devcel.2020.11.013.

34. Pijuan-Sala, B., Griffiths, J.A., Guibentif, C., Hiscock, T.W., Jawaid, W., Calero-Nieto, F.J., Mulas, C., Ibarra-Soria, X., Tyser, R.C.V., Ho, D.L.L., et al. (2019). A single-cell molecular map of mouse gastrulation and early organogenesis. Nature 566, 490–495. 10.1038/s41586-019-0933-9.

35. Hamburger, V., and Hamilton, H.L. (1992). A series of normal stages in the development of the chick embryo (1951). Dev.Dyn. 195, 231–272.

36. Henrique, D., Abranches, E., Verrier, L., and Storey, K.G. (2015). Neuromesodermal progenitors and the making of the spinal cord. Development 142, 2864–2875. 10.1242/dev.119768.

37. Furtado, M.B., Solloway, M.J., Jones, V.J., Costa, M.W., Biben, C., Wolstein, O., Preis, J.I., Sparrow, D.B., Saga, Y., Dunwoodie, S.L., et al. (2008). BMP/SMAD1 signaling sets a threshold for the left/right pathway in lateral plate mesoderm and limits availability of SMAD4. Genes Dev 22, 3037–3049. 10.1101/gad.1682108.

38. Schiebinger, G., Shu, J., Tabaka, M., Cleary, B., Subramanian, V., Solomon, A., Gould, J., Liu, S., Lin, S., Berube, P., et al. (2019). Optimal-Transport Analysis of Single-Cell Gene Expression Identifies Developmental Trajectories in Reprogramming. Cell 176, 928–943 e922. 10.1016/j.cell.2019.01.006.

39. Loulier, K., Barry, R., Mahou, P., Le Franc, Y., Supatto, W., Matho, K.S., Ieng, S., Fouquet, S., Dupin, E., Benosman, R., et al. (2014). Multiplex cell and lineage tracking with combinatorial labels. Neuron 81, 505–520. 10.1016/j.neuron.2013.12.016.

40. Serra, M., Streichan, S., Chuai, M., Weijer, C.J., and Mahadevan, L. (2020). Dynamic morphoskeletons in development. Proc Natl Acad Sci U S A 117, 11444–11449. 10.1073/pnas.1908803117.

41. Benazeraf, B., Beaupeux, M., Tchernookov, M., Wallingford, A., Salisbury, T., Shirtz, A., Shirtz, A., Huss, D., Pourquie, O., Francois, P., and Lansford, R. (2017). Multi-scale quantification of tissue behavior during amniote embryo axis elongation. Development. 10.1242/dev.150557.

42. Moreau, C., Caldarelli, P., Rocancourt, D., Roussel, J., Denans, N., Pourquie, O., and Gros, J. (2019). Timed Collinear Activation of Hox Genes during Gastrulation Controls the Avian Forelimb Position. Curr Biol 29, 35–50 e34. 10.1016/j.cub.2018.11.009.

43. Vermillion, K.L., Bacher, R., Tannenbaum, A.P., Swanson, S., Jiang, P., Chu, L.F., Stewart, R., Thomson, J.A., and Vereide, D.T. (2018). Spatial patterns of gene expression are unveiled in the chick primitive streak by ordering single-cell transcriptomes. Dev Biol 439, 30–41. 10.1016/j.ydbio.2018.04.007.

44. Qiu, C., Cao, J., Martin, B.K., Li, T., Welsh, I.C., Srivatsan, S., Huang, X., Calderon, D., Noble, W.S., Disteche, C.M., et al. (2022). Systematic reconstruction of cellular trajectories across mouse embryogenesis. Nat Genet 54, 328–341. 10.1038/s41588-022-01018-x.

45. Qiu, C., Martin, B.K., Welsh, I.C., Daza, R.M., Le, T.M., Huang, X., Nichols, E.K., Taylor, M.L., Fulton, O., O’Day, D.R., et al. (2024). A single-cell time-lapse of mouse prenatal development from gastrula to birth. Nature 626, 1084–1093. 10.1038/s41586-024-07069-w.

46. Ton, M.N., Keitley, D., Theeuwes, B., Guibentif, C., Ahnfelt-Ronne, J., Andreassen, T.K., Calero-Nieto, F.J., Imaz-Rosshandler, I., Pijuan-Sala, B., Nichols, J., et al. (2023). An atlas of rabbit development as a model for single-cell comparative genomics. Nat Cell Biol 25, 1061–1072. 10.1038/s41556-023-01174-0.

47. VanHorn, S., and Morris, S.A. (2021). Next-Generation Lineage Tracing and Fate Mapping to Interrogate Development. Dev Cell 56, 7–21. 10.1016/j.devcel.2020.10.021.

48. Fulton, T., Verd, B., and Steventon, B. (2022). The unappreciated generative role of cell movements in pattern formation. R Soc Open Sci 9, 211293. 10.1098/rsos.211293.

49. Firmino, J., Rocancourt, D., Saadaoui, M., Moreau, C., and Gros, J. (2016). Cell Division Drives Epithelial Cell Rearrangements during Gastrulation in Chick. Dev Cell 36, 249–261. 10.1016/j.devcel.2016.01.007.

50. Hayward, M.K., Muncie, J.M., and Weaver, V.M. (2021). Tissue mechanics in stem cell fate, development, and cancer. Dev Cell 56, 1833–1847. 10.1016/j.devcel.2021.05.011.

51. McGrew, M.J., Sherman, A., Ellard, F.M., Lillico, S.G., Gilhooley, H.J., Kingsman, A.J., Mitrophanous, K.A., and Sang, H. (2004). Efficient production of germline transgenic chickens using lentiviral vectors. EMBO Rep 5, 728–733.

52. Hamburger, V. (1992). The stage series of the chick embryo [comment]. Dev Dyn. 195, 273–275.

53. Zacchei, A.M. (1961). [The embryonal development of the Japanese quail (Coturnix coturnix japonica T. and S.)]. Arch Ital Anat Embriol 66, 36–62.

54. Chapman, S.C., Collignon, J., Schoenwolf, G.C., and Lumsden, A. (2001). Improved method for chick whole-embryo culture using a filter paper carrier. Dev Dyn 220, 284–289.

55. Oginuma, M., Moncuquet, P., Xiong, F., Karoly, E., Chal, J., Guevorkian, K., and Pourquie, O. (2017). A Gradient of Glycolytic Activity Coordinates FGF and Wnt Signaling during Elongation of the Body Axis in Amniote Embryos. Dev Cell 40, 342–353 e310. 10.1016/j.devcel.2017.02.001.

56. Hama, H., Kurokawa, H., Kawano, H., Ando, R., Shimogori, T., Noda, H., Fukami, K., Sakaue-Sawano, A., and Miyawaki, A. (2011). Scale: a chemical approach for fluorescence imaging and reconstruction of transparent mouse brain. Nat Neurosci 14, 1481–1488. 10.1038/nn.2928.

